# Vermeer: Autoregressive generative modeling of microscopy predicts protein localization

**DOI:** 10.64898/2026.06.01.729395

**Authors:** Sandeep Kambhampati, Eric Zimmermann, Emre Hayir, Kevin K. Yang, Fei Chen, Alex X. Lu

## Abstract

Fluorescent microscopy provides a rich view into how proteins localize within cells, but it remains experimentally infeasible to image human proteins across all of the different factors that can impact localization. We introduce *Vermeer*, a channel-adaptive autoregressive generative model for *in silico* generation of microscopy images of protein localization. Vermeer conditions generations on protein sequences and landmark stains showing the morphology of cells, which enables it to generalize to unseen proteins and cell lines. We show that Vermeer, trained on the Human Protein Atlas, can generate images with substantially improved perceptual quality and biological fidelity over previous proposals. Additionally, Vermeer’s autoregressive framework enables flexible generation using varying channel subsets and orderings, enabling zero-shot transfer to data collected under different imaging conditions and channel configurations than those used for training. These results position Vermeer to enable scalable modeling of protein localization and are a step towards generative foundation models that can operate over distinct microscopy datasets. Code is publicly accessible at https://github.com/microsoft/vermeer.git.

## 1 Introduction

Proteins drive nearly all cellular processes, but their function is dependent on localizing to the correct cellular component [1]. Mislocalization due to mutations is associated with many diseases [2]. Importantly, protein localization is not determined by amino acid sequence alone: the same protein can exhibit distinct localization patterns across cell types, cellular states, and environmental conditions, often reflecting differences in biological function [1]. For this reason, even though a protein’s amino acid sequence contains information about where it *may* localize, experimentation is essential for determining where proteins actually localize in response to both intrinsic cellular state and external environmental cues [1]. Fluorescence microscopy provides a powerful approach for directly visualizing protein localization, enabling the development of large-scale imaging resources that systematically characterize the subcellular distribution of proteins [2, 3, 4, 5, 6, 7].

However, comprehensive experimental data collection remains intractable. For example, the Human Protein Atlas has collected images for 13,080 human proteins across 40 different cell lines [3]. Because of the immense labor and expense involved, only 8% of all protein and cell line combinations have been collected to date. Even then, the Human Protein Atlas only images 65% of all human proteins, and just 40 out of thousands of possible cell lines. As cell line is not the only factor that will impact protein localization, considering other factors such as environmental perturbations or genetic mutations in conjunction means the space of experimental conditions grows combinatorially.

Recent advances in generative modeling offer a promising approach to addressing this challenge by synthesizing images of fluorescently labeled proteins [8, 9, 10]. These models are typically conditioned on a protein sequence and base microscopy images showing “landmark stains” visualizing the morphology of a specified cell. Because these generative models can predict how different proteins will localize in cells that have *not* been stained with the protein, they can prioritize more critical proteins to target for experimentation (e.g. those that are differentially localized compared to a baseline condition). Importantly, generating spatially resolved images offers advantages over predicting discrete subcellular compartment labels. Protein localization often exhibits complex spatial patterns, including heterogeneous distributions, punctate structures, and gradients, that may distinguish functional states even within the same organelle. These features contain biological information that is lost when localization is reduced to a categorical annotation [11, 12, 13].

However, many existing methods are unable to leverage this advantage of image generation, because they generate images that show fuzzy distributions over the cell instead of precisely localizing proteins. This firstly limits the resolution of biological function, as functionally different organelles can exhibit similar spatial distributions in a 2D image, and can only be distinguished by sharper features like texture. This also means that existing models do not faithfully reproduce localization cues like intense foci or gradients that point to biological function. Furthermore, all existing models assume fixed channel input. However, different microscopy datasets may have different numbers of channels depending on experimental setup. This means that models need to be explicitly re-trained for each new dataset they encounter.

Here, we introduce Vermeer, an autoregressive generative model trained on the Human Protein Atlas. We advance several key contributions:

- Trained on the Human Protein Atlas, Vermeer generates images that accurately capture fine-grained textural and localization cues across different organelles, scoring higher than previous methods on measures of perceptual quality and biological plausibility. We also demonstrate Vermeer can generalize zero-shot to image resolutions unseen in training, enabling transfer to datasets with varying imaging conditions.
- We validate that Vermeer can more accurately generate images of protein localization for unseen cell lines of different tissue type and unseen proteins. To explain this generalization, we show Vermeer systematically detects localization signals in protein sequences.
- Our autoregressive framework means that Vermeer can generate images of protein localization for arbitrary numbers of landmark stain channels, making it channel-adaptive. We exploit this property to demonstrate that Vermeer can generalize zero-shot to out-of-distribution datasets with different numbers of channels as training.

## 2 Related Works

### 2.1 Generative Image Models for Protein Localization

Several existing generative models aim to predict the subcellular localization of proteins by generating images conditionally given landmark stains and protein sequences. PUPS [8] trains a U-Net decoder style model, minimizing a mean-squared error loss on generated pixels. CELL-E 2 [9] trains a bi-directional transformer, minimizing a cross-entropy loss on generated tokens. However, both models produce low-quality generated images that cannot resolve localization between many organelles. CELL-Diff [10] trains a latent diffusion model producing higher-quality generations. Here, we further extend by exploring autoregressive vision models, which unlocks new applications for cross-dataset generalization.

For these models to be useful, they must generalize to unseen protein sequences and cell lines. CELL-E 2 only evaluates on unseen protein sequences. PUPS evaluates on unseen proteins and cell lines, which we also follow, but only evaluates generations on low-level pixel statistics (e.g. mean-squared error) or on distribution at the coarsest cellular resolution (cytoplasm-nuclear ratio). We additionally verify the biological fidelity of generated images by using a localization classifier which predicts labels over 31 localization classes and comparing to ground-truth annotations. We also test zero-shot generalization to unseen image resolutions, and to new datasets with distinct staining procedures, which no previous work tests.

### 2.2 Channel Adaptive Microscopy Methods

Different microscopy datasets may have different numbers of channels depending on experimental setup. This variability poses a challenge for vision models that assume a fixed number of input channels. Recent work for microscopy has therefore explored channel-adaptive architectures for representation learning, including ChAdaViT [14], CA-MAE [15], and ChA-MAEViT [16]. To date, channel-adaptive models have not been applied for generating microscopy images. Here, we demonstrate the value of channel adaptivity for generation by showing zero-shot generalization to a dataset with a different number of landmark channels than training. We further extend channel-adaptive models to incorporate information about protein sequences and enable conditional generation of protein localization.

## 3 Methods

### 3.1 Overview

Vermeer extends autoregressive image modeling for natural images to multi-channel microscopy images (Figure 1). In contrast to natural images where RGB channels encode correlated color information, each channel of a microscopy image is a unique biological marker. Following previous channel-adaptive modeling proposals [14, 15, 16], we therefore treat each microscopy channel as an independent input. The model follows a two-stage pipeline: (1) each channel of a microscopy image is independently tokenized into a discrete code sequence using a pretrained VQ-VAE encoder [17]; (2) a transformer is trained to autoregressively generate a single sequence formed by concatenating the per-channel token sequences. Generated tokens can then be decoded by the VQ-VAE back into pixels.

**Figure 1:**
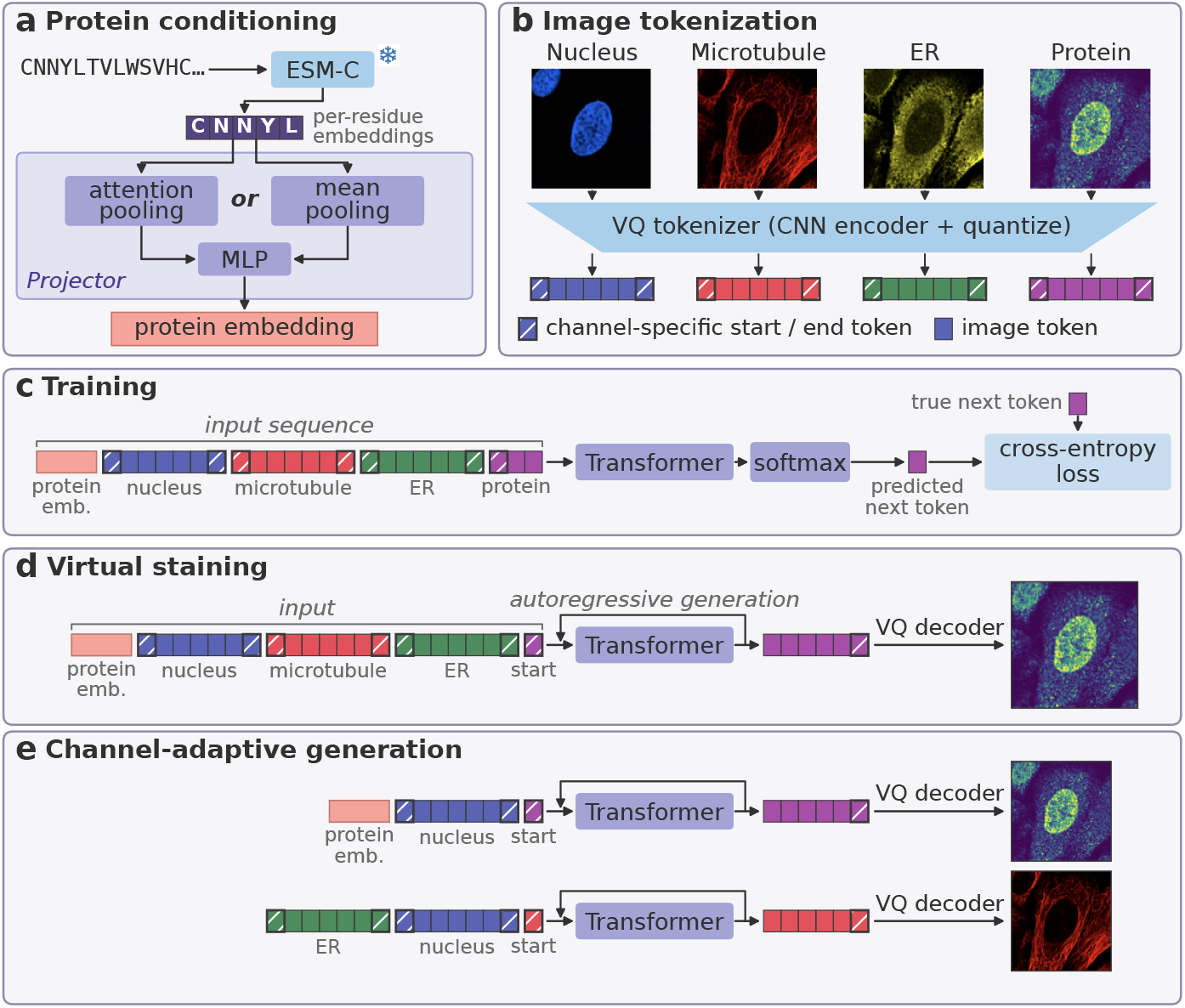
Vermeer model overview. Vermeer is a channel-adaptive autoregressive model for fluorescent microscopy data. (a) Vermeer can take in conditioning input, such as protein sequence embeddings. Protein sequence embeddings are pooled over amino acids via either attention pooling or mean pooling. (b) Vermeer tokenizes each input channel independently via a patch-wise raster-scan order and adds channel-specific delimiter (“start / end”) tokens. (c) The model is trained on a next-token prediction task across all channels. “+” indicates a concatenation operation. (d) At inference time, Vermeer can generate a protein stain conditioned on input protein sequence and reference stains of cell morphology. (e) We additionally train a variant, Vermeer-CA, that permutes and drops out channels during training. At inference time, this allows Vermeer-CA to be flexible to the specific order and subset of channels and can be prompted to generate any channel it has seen during training.

Training can be viewed as sequential generation across channels: after the first channel is generated subsequent channels are generated conditionally, with the generated channel *c* conditioned on the preceding *c* ™ 1 channels. We refer to conditional generation on other preceding channels as “virtual staining” as short-hand. This formulation encourages the model to capture cross-channel dependencies by learning the joint distribution over channel-specific token sequences. We additionally incorporate a prefix embedding that serves as conditioning information for the generations. In the case of our model, this prefix is the ESM-C [18, 19] embedding of protein sequence.

The backbone is a decoder-only transformer following the Llama architecture [20, 21], as adapted by LlamaGen [17]. We report results for a 775M parameter model (Vermeer-XL), but also train architectures at two other scales (Table S1). We find that model scaling improves performance on downstream evaluation metrics (Table S2, S3, S4). We additionally train two variants of models: our base Vermeer models, which use a fixed channel order, and Vermeer-CA, which randomly permutes channel order and drops out channels during training (Figure 1E).

### 3.2 Multi-channel Image Tokenization

We adopt the tokenizer architecture from LlamaGen [17], which builds on Esser et al. [22]. The encoder maps a 256 × 256 grayscale image (replicated to 3 channels) to a 16 × 16 latent grid via 16× spatial downsampling. Therefore, each token approximately corresponds to a 16 pixel x 16 pixel patch in input space. Each spatial position is quantized to the nearest entry in a learned codebook of size |*V*| = 16,384, yielding *P* = 256 discrete tokens per channel. We use VQ-VAE weights pretrained on ImageNet [17] for tokenization.

To model multi-channel microscopy images as a single autoregressive sequence, we “flatten” the image across the channel dimension. We augment the base VQ codebook *V* = {1, …, |*V*|} with channel-specific delimiter tokens and a sequence terminator. For a maximum of *C*_max_ channels, the extended vocabulary V_ext_ is:

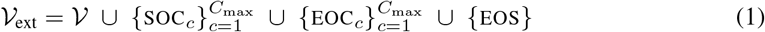

A multi-channel image with *C*_*max*_ channels is therefore represented as:

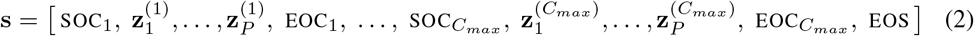

where 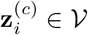 is the *i*-th VQ code for channel *c*. The delimiter tokens provide an explicit signal of channel identity and enable flexible prompting for the generation of specific channels at inference time, as well as varying channel order for Vermeer-CA specifically. All images share the same delimiter token for the protein channel. We additionally use 2D rotary positional embeddings (RoPE) [23] to encode spatial positions of image patches, using the same spatial coordinate system across channels.

### 3.3 Protein Sequence Conditioning

We condition generation on protein sequence information extracted by ESM-C 600M [18], a protein language model that produces per-residue embeddings of dimension *d*_ESM_ = 1152. Given a protein sequence of length *L*, we extract embeddings 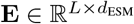 from ESM-C. We investigate two pooling strategies (Figure 1A) to produce a fixed-size conditioning vector. For both strategies, the pooled embedding is projected to the model dimension via a two-layer MLP with GELU activation. Unless otherwise mentioned, results are reported for the model using mean-pooling.

#### Mean-pooling

The simplest approach averages over residue positions *i*:

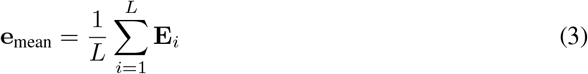

#### Attention pooling

To enable the model to focus on informative regions of the sequence, we employ an attention-based pooling mechanism inspired by DeepLoc 2.0 [24]. Given per-residue embeddings 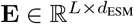 and a learnable query vector 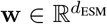:

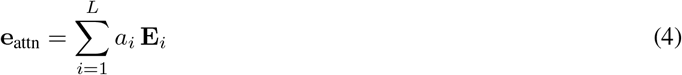

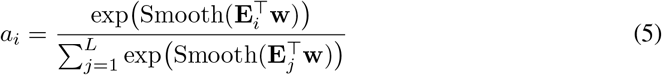

where Smooth(·) refers to a smoothing operation with a Gaussian filter (*σ* = 2). This incorporates the inductive bias that sequence motifs exist in a contiguous set of residues. The learned, smoothed attention weights *a*_*i*_ can subsequently be interpreted to identify informative regions of the protein sequence.

### 3.4 Training Objective

We minimize a standard autoregressive cross-entropy loss:

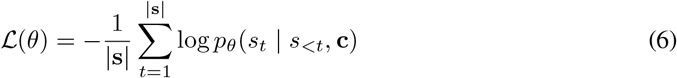

where **c** is the conditioning embedding and *θ* denotes model parameters. Further details on training and inference can be found in Appendix A.1–A.6.

### 3.5 Datasets

#### Human Protein Atlas

The Human Protein Atlas [3] is a multi-channel immunofluorescence dataset profiling 13,080 proteins (65% of the human proteome). Most proteins are profiled in three cell lines, across 40 total cell lines. The dataset consists of 95K images, each with 4 channels: nucleus, microtubule, endoplasmic reticulum (ER), and a protein of interest. The training and validation sets are designed to evaluate generalization across both proteins and cell lines (Appendix A.7). Specifically, we construct three types of validation splits (Figure S1): (1) an unseen protein × cell line combination split, where specific protein–cell line pairs are excluded during training; (2) an unseen protein split, where entire proteins are held out across all cell lines; and (3) an unseen cell-line split, where entire cell lines are excluded during training. The unseen cell-lines are chosen to test increasing degrees of generalization, including cell-types and organs not seen during training (Appendix A.7).

#### OpenCell

OpenCell [25] is a live-cell imaging dataset using CRISPR-based fluorescent labeling to endogenously tag proteins. The OpenCell database consists of 6,301 images, each with 2 channels: nucleus and a protein of interest. We use OpenCell for out-of-distribution validation, as it differs from the HPA in its channel composition and data distribution (1124 proteins overlap while 187 proteins are unique), making it a stringent test of robustness across both biological and technical domain shifts.

### 3.6 Evaluation

We compute several metrics to assess whether the generated images are consistent with the real images. We compute the Frechet Inception Distance (FID) [26] as well as the pixel-level mean-squared error (MSE). For comparison with PUPS, we measure their proposed metric of whether nuclear vs cytoplasmic localization is correctly generated by computing the Pearson Correlation Coefficient (PCC) between the intra-nuclear proportion calculated from real and generated images.

To evaluate whether generated protein channel images capture biologically meaningful patterns, we pass both real and generated images through SubCellPortable, a protein subcellular localization classifier trained on real HPA images [4]. This oracle produces probability distributions over 31 localization classes (e.g., nucleoplasm, mitochondria, cytosol). The oracle achieves close to stateof-the-art performance on classifying localization of images from the Human Protein Atlas [4]. We compare predictions on generated images against (a) predictions on real images and (b) ground-truth HPA annotations. We report both the Spearman Correlation between the vector of probabilities assigned to each class for the true image versus the generated image, along with additional metrics that capture the accuracy of the prediction relative to ground-truth HPA annotations. Further details on the specific metrics used can be found in Appendix A.8.

## 4 Results

### 4.1 Vermeer generates more biologically plausible images than previous proposals

Existing image databases do not profile all proteins in all cell-lines, due to the expense and labor in systematically scaling experimentation. We therefore first evaluate our model on unseen protein × cell-line pairs, where both the protein and cell-line may be observed during training but not in that specific combination. Qualitatively, we observe Vermeer generates images with higher perceptual quality than existing methods (Figure S2, S3). In contrast, previous methods such as PUPS [8] and CELL-E 2 [9] generate blurry and unrealistic images.

To complement qualitative results, we quantified image generation on metrics of both perceptual quality and biological plausibility (Table 1).We find that Vermeer outperforms all prior models on all metrics. We further compare Vermeer against naive strategies of randomly scrambling pixels (which preserves intensity distribution but not localization), and using Vermeer but with a random protein prefix (which verifies correct generation of protein localization depends upon protein sequence). We verify that Vermeer outperforms these naive baselines as well. Although we report the results for our largest model, we also observe that model scaling improves performance on evaluation metrics (Figure S4, S5).

**Table 1:** Benchmarking results on unseen protein × cell line split. SubCell classification metrics score SubCell predictions of localization class from the generated image against annotated labels. Best metrics are bolded.

| Model | FID<br>(↓) | MSE<br>(↓) | PCC<br>Intranuc. Prop.<br>(↑) | SubCell<br>Spearman Corr.<br>(↑) | SubCell Classification Metrics |  |  |  |
|---|---|---|---|---|---|---|---|---|
|  |  |  |  |  | Coverage Error<br>(↓) | MLRAP<br>(↑) | Macro AP<br>(↑) | Micro AP<br>(↑) |
| CELL-E 2 | 286 | 0.2213 | 0.246 | 0.650 | 7.70 | 0.454 | 0.062 | 0.243 |
| CELL-Diff | 42.1 | 0.0586 | 0.329 | 0.700 | 7.66 | 0.495 | 0.096 | 0.294 |
| PUPS | 102 | 0.0534 | 0.828 | 0.855 | 7.49 | 0.610 | 0.172 | 0.477 |
| Vermeer-XL | **15.6** | **0.0373** | **0.884** | **0.856** | **5.28** | **0.662** | **0.190** | **0.526** |
| *Vermeer-XL baselines* |  |  |  |  |  |  |  |  |
| Pixel shuffled | 459 | 0.0929 | 0.182 | 0.559 | 8.47 | 0.380 | 0.065 | 0.203 |
| Protein prefix shuffled | 17.2 | 0.0583 | 0.169 | 0.745 | 8.00 | 0.460 | 0.059 | 0.254 |

One challenge to systematic benchmarking is that all these previous models operate on different microscopy resolutions, preprocessing, and data splits. We test how much these factors may alter the ranking of model performance in two more controlled setups. First, we train all baselines on our data split and processing, at our resolution. We verify that Vermeer outperforms on all metrics in this setting (Table S5). Second, we evaluate Vermeer zero-shot at resolutions unseen during training (Table S5). Surprisingly, Vermeer still outperforms all other methods, even when applied to image resolutions out-of-distribution from what was used for training. These results indicate that Vermeer has capacity to generalize to image resolutions unseen during training zero-shot. Further details on the benchmarking protocol can be found in Appendix A.9.

We verified that Vermeer’s generations do not rely on spectral bleedthrough — a phenomenon in fluorescent microscopy where faint signal from one channel leaks into other imaging channels. Prior work (PUPS) showed that naive models can overfit to this artifact, and thereby access information about the protein stain from the provided landmark stains, and they mitigate by training models on strongly thresholded data.

Following PUPS, we generated proteins not present in the original stains while keeping landmark channels fixed. The resulting localizations match the prompted protein identities rather than the ground-truth stains (Figure S6), indicating that Vermeer does not depend on bleedthrough. Unlike PUPS, this emerges with no special thresholding of images. We speculate this is because our frozen pretrained VQ-VAE tokenizer does not capture this faint signal.

### 4.2 Vermeer generalizes to unseen proteins and unseen cell-lines

Existing image data collections only profile a subset of the human proteome. A major reason for this limited coverage is technical limitations on antibody validation [3] or GFP-tagging [25], therefore placing many proteins outside of current experimental capabilities. To assess if Vermeer can generate proteins outside of those currently experimentally viable for imaging, we test the ability of the model to generalize to new proteins outside of the training data.

We indeed find that Vermeer exhibits capacity to generalize to unseen proteins (Figure 2), and the model is able to generate accurate protein stains across localization classes. The model outperforms all existing proposals on FID, MSE, and SubCell classification metrics, while PUPS outperforms on PCC Intranuclear Proportion and SubCell Spearman Correlation (Table 2, Table S6). This suggests that there may be some trade-offs between Vermeer and PUPS, where Vermeer may better generate the fine-grained localization class at the expense of accuracy of relative distribution between the nucleus and cytoplasm for unseen proteins specifically.

**Table 2:** Benchmarking results on unseen protein split. SubCell classification metrics score SubCell predictions of localization class from the generated image against annotated labels. Best metrics are bolded.

| Model | FID<br>(↓) | MSE<br>(↓) | PCC<br>Intranuc. Prop.<br>(↑) | SubCell<br>Spearman Corr.<br>(↑) | SubCell Classification Metrics |  |  |  |
|---|---|---|---|---|---|---|---|---|
|  |  |  |  |  | Coverage Error<br>(↓) | MLRAP<br>(↑) | Macro AP<br>(↑) | Micro AP<br>(↑) |
| CELL-E 2 | 287 | 0.2232 | 0.265 | 0.648 | 7.48 | 0.466 | 0.065 | 0.247 |
| CELL-Diff | 41.9 | 0.0589 | 0.340 | 0.699 | 7.45 | 0.506 | 0.092 | 0.307 |
| PUPS | 104 | 0.0691 | **0.652** | **0.825** | 8.28 | 0.538 | 0.092 | 0.369 |
| Vermeer-XL | **18.1** | **0.0470** | 0.624 | 0.802 | **6.48** | **0.576** | **0.113** | **0.399** |
| *Vermeer-XL baselines* |  |  |  |  |  |  |  |  |
| Pixel shuffled | 457 | 0.0921 | 0.171 | 0.560 | 8.16 | 0.390 | 0.072 | 0.209 |
| Protein prefix shuffled | 19.5 | 0.0554 | 0.225 | 0.759 | 7.61 | 0.491 | 0.065 | 0.284 |

**Figure 2:**
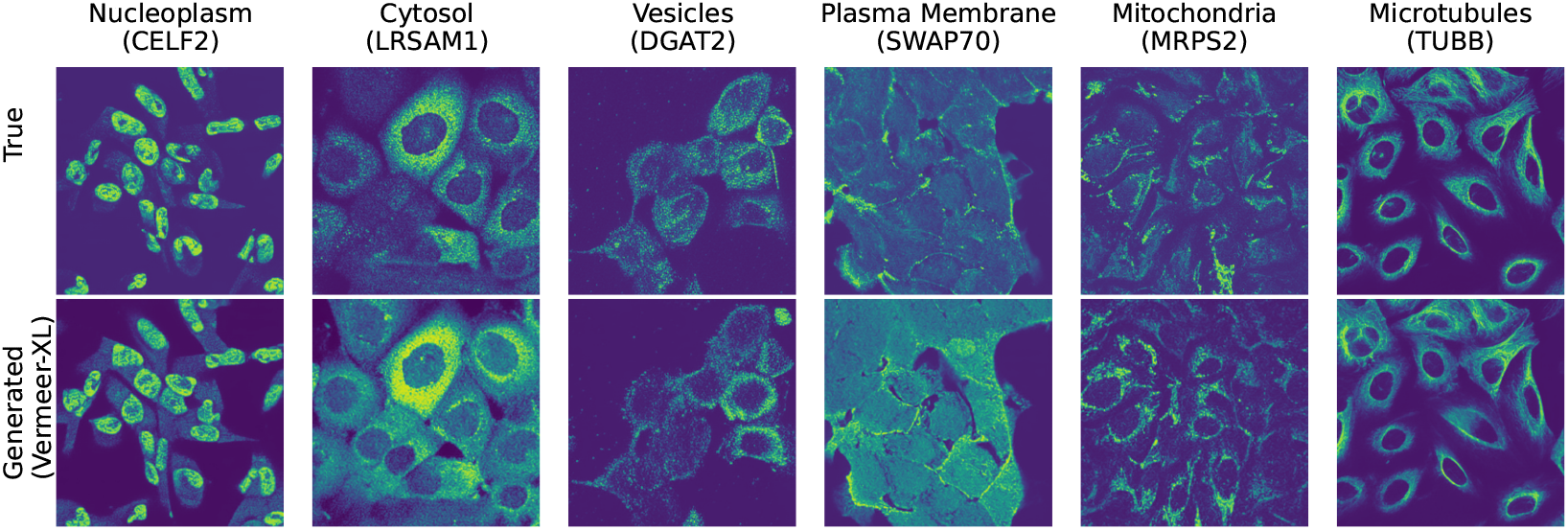
Example protein virtual staining results on unseen proteins across localization classes.

We evaluate the degree to which Vermeer relies on the protein language model embeddings to generalize to unseen proteins. To do this, we train a baseline that replaces ESM embeddings with a learnable embedding for each training protein. For an unseen protein, we find its nearest training protein in ESM embedding space using cosine similarity and use that protein’s learned embedding. We find that on the unseen protein split, this stringent memorization baseline performs worse than conditioning directly on the ESM embedding (Table S7). This suggests that Vermeer is successfully leveraging latent information regarding protein localization in the ESM embedding space to make its predictions.

We do note a gap in performance on evaluation metrics compared to proteins seen during training. We find that performance is moderately correlated with the sequence similarity of unseen proteins to proteins in the training set, but less correlated with the cosine similarity of ESM embeddings between unseen proteins and train proteins. (Figure S7). We also find that performance is correlated with the frequency of the localization class in training. We note that the gap in performance between seen and unseen proteins is largely driven by the rarer localization classes (Figure S8). The model achieves comparable performance on both unseen and seen proteins for the two most common classes in the data, nuclear and cytoplasmic.

We also analyze how localization variability affects model performance by stratifying proteins into three categories (Appendix A.10). We observe that increased localization variability is inversely correlated with performance for both seen and unseen proteins (Figure S9). This suggests that the model may not capture the full extent of biological heterogeneity in protein localization.

We further test whether Vermeer learns meaningful relationships between cell morphology and subcellular localization that may enable the model to generalize to unseen cell-lines. We find that Vermeer is indeed able to generate protein stains for unseen cell-lines with high biological fidelity, with comparable performance across individual cell-lines (Figure S10). The oracle classifier’s performance on generated protein stains for unseen cell-lines approaches that of the true protein stains (Table S4).

### 4.3 Vermeer attends to parts of protein sequence driving localization

As Vermeer shows some generalization to unseen protein sequences, we next sought to understand the basis of this generalization. To do this, we use a variant of the Vermeer model that applies attention pooling across individual amino acid residues, and assess if the model puts higher attention on localization-driving parts of the protein sequence.

We focus on three examples of localization subsequences: nuclear localization signals, endoplasmic reticulum (ER) signal peptides, and mitochondrial transit peptides, which regulate localization into the nucleus, ER, and mitochondria, respectively. We use annotations of known localization subsequences from Uniprot [27] and compute the attention weights on residues that overlap with these known subsequences and the residues that do not. We find that across all three localization signals, the model has higher attention weights on residues that overlap with a known subsequence (Figure 3) than the rest of the residues in the same proteins. Together, these results suggest that Vermeer is able to identify and attend to biologically meaningful sequence features associated with subcellular localization.

**Figure 3:**
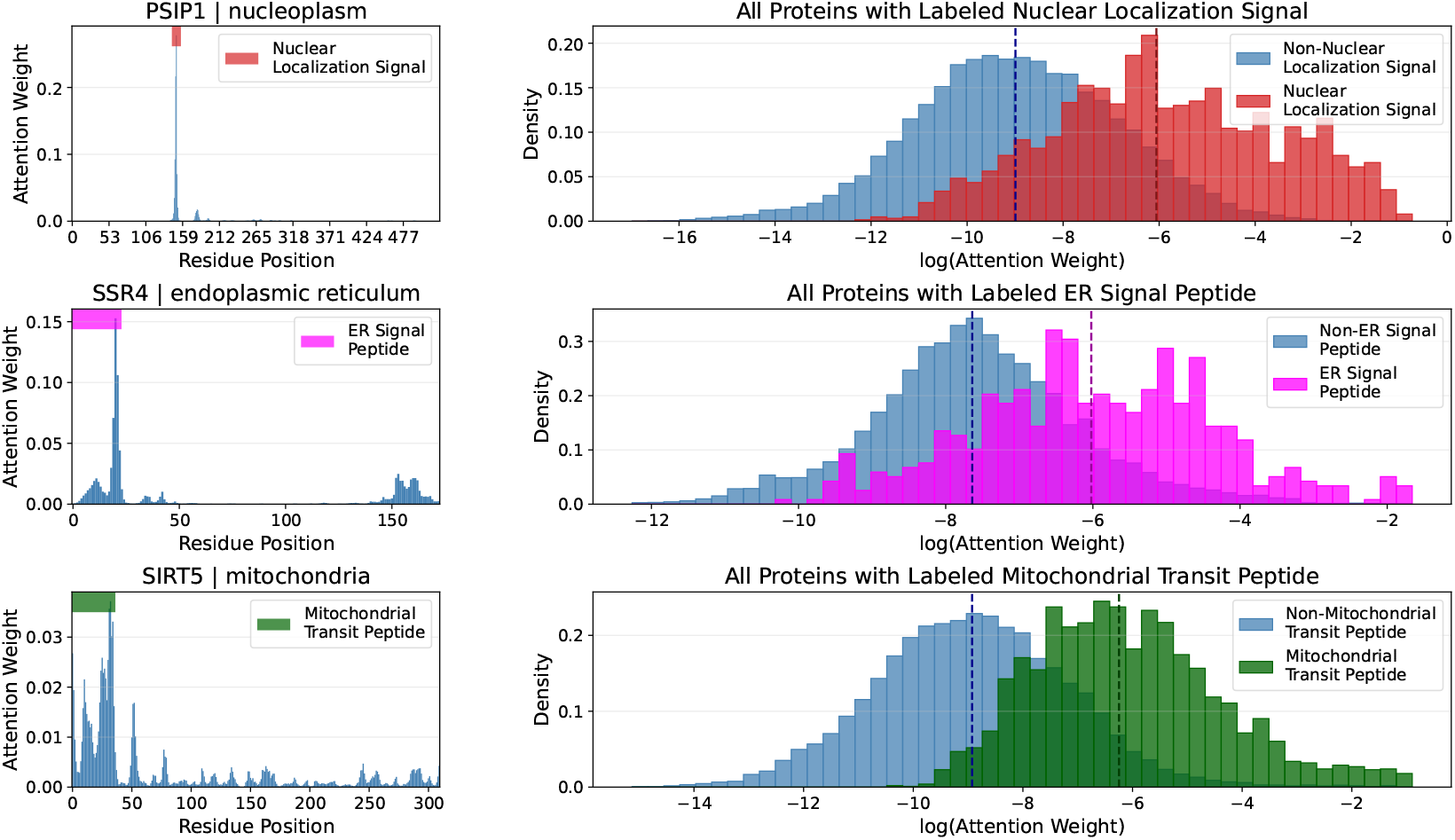
Interpretability analysis of the learned attention pooling mechanism with Vermeer-AP. We show the distribution of attention values on known localization subsequences, for amino acid residues belonging to the subsequence versus other residues in the protein.

### 4.4 Vermeer enables channel-adaptive generations

Next, to demonstrate the flexibility of our autoregressive approach, we trained a variant of our model (Vermeer-CA) that randomly permutes the order of channels. We reasoned this would enable Vermeer to generalize during inference to new datasets, which may collect some overlapping landmark stains, but not the exact same set of landmark stains used during training. By permuting channels during training, this means sometimes the nucleus channel is the first channel, and the protein channel directly follows, enabling generation of proteins of interest given a nucleus landmark stain alone.

We first demonstrate, qualitatively, Vermeer-CA can still perform protein virtual staining given only a subset of the landmark stains for held-out images within the Human Protein Atlas (Figure S11). We next tested protein virtual staining on an unseen dataset, OpenCell [25], which contains live-cell images with only a nuclear channel and GFP-tagged proteins. Despite differences in resolution, labeling, and channels, the model generates plausible localizations (Figure 4). However, these do not always match the true protein patterns, possibly due to biological covariate shifts between Human Protein Atlas and OpenCell. In particular, GFP-tagging and live-cell imaging can alter localization patterns compared to antibody-based staining. Thus, while Vermeer-CA addresses technical generalization challenges, it remains unclear if it fully transfers biologically. We also note that Vermeer-CA slightly underperforms Vermeer-XL, suggesting that generalization to datasets with different landmark stain channels may come at expense of some performance on the original dataset of interest.

**Figure 4:**
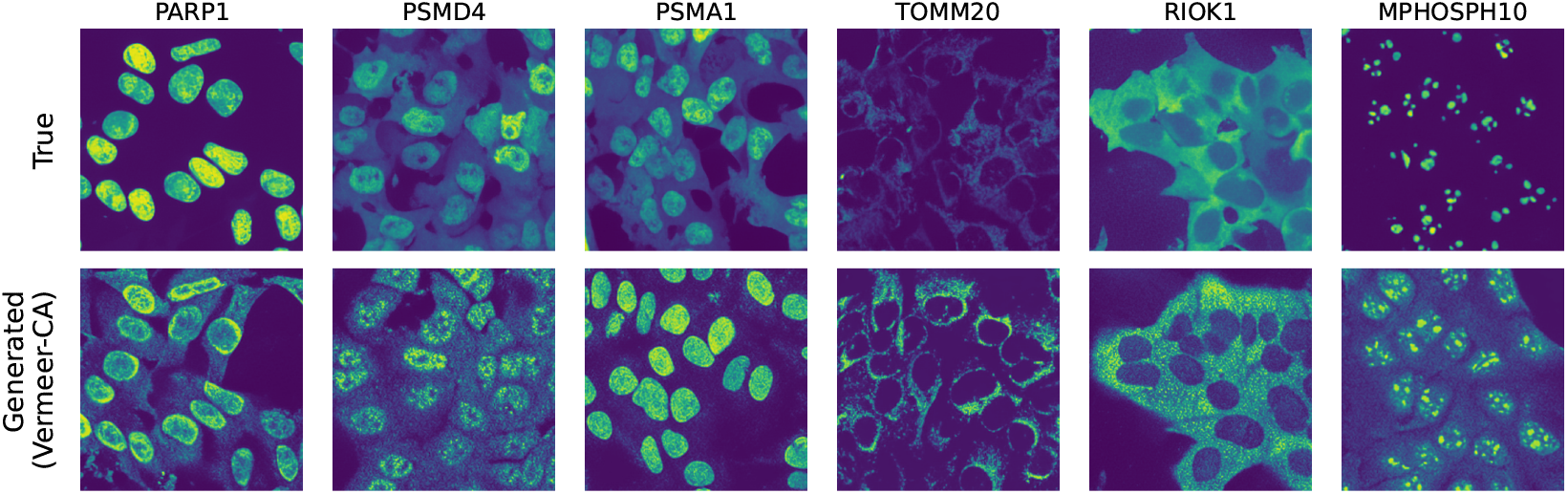
Example protein virtual staining results from zero-shot transfer to the OpenCell dataset

We also compare our channel-adaptive approach to a naive adaptation of a fixed-channel model, CELL-Diff, where missing landmark stain channels in a new dataset are filled with blank images. We run this comparison across two settings: first, in the zero-shot setting, where models are only trained on the Human Protein Atlas data, and second, after fine-tuning on the OpenCell data. We find that Vermeer-CA significantly outperforms this approach across all image generation metrics (Table 3), in both cases.

**Table 3:** Image generation performance on OpenCell.

| Model | FID (↓) | MSE (↓) | PCC Intranuc Prop (↑) |
|---|---|---|---|
| Vermeer-CA (zero-shot) | **71.1** | **0.0545** | **0.592** |
| CELL-Diff (zero-shot) | 498 | 0.269 | 0.430 |
| Vermeer-L (finetuned) | **7.47** | **0.0217** | **0.928** |
| CELL-Diff (finetuned) | 51.6 | 0.0392 | 0.798 |

## 5 Conclusion

Vermeer provides a scalable and flexible framework for the generation of fluorescent microscopy data. By modeling protein sequence embeddings and image tokens within a unified autoregressive framework, Vermeer enables biologically grounded generation of protein localization patterns while remaining flexible to varying channel subsets and orderings. We establish rigorous validation settings to test generalization to unseen protein × cell-line combinations, entirely unseen proteins, and entirely unseen cell-lines, and show that Vermeer produces realistic and biologically meaningful generations across these settings. Together, these results suggest that autoregressive, channel-adaptive modeling is a promising approach for building generative foundation models for heterogeneous microscopy data.

However, several challenges remain. Performance degrades on rare and highly variable localization patterns, highlighting the difficulty of fully capturing biological heterogeneity. The Human Protein Atlas is also relatively limited in size, and future work could leverage Vermeer’s channel-adaptive capabilities to train on larger and more diverse microscopy datasets [28].

More broadly, Vermeer provides a foundation for learning the relationship between protein sequence, cellular context, and subcellular localization. Vermeer can be readily extended to be natively multimodal across protein sequence and image tokens, and thus can provide a foundation for future *de novo* protein design tasks, such as generating proteins with desired localization patterns. By enabling computational predictions of protein localization across conditions that would be impractical to characterize experimentally, Vermeer offers a path toward scalable exploration of protein behavior and the discovery of previously uncharacterized mechanisms of subcellular organization.

## 6 Acknowledgements

We thank Dinghuai Zhang, David Alvarez-Melis, and Shantanu Singh for helpful discussions during the course of this project. We are also grateful to members of Microsoft Research New England and the Chen Lab for their support, feedback, and guidance.

## 7 Code and Data Availability

Code is publicly accessible at https://github.com/microsoft/vermeer.git. Instructions on downloading the full Human Protein Atlas dataset, as well as a toy example and tutorial notebook are provided in the Github README. Vermeer model weights are publicly accessible from HuggingFace (see README for details). The OpenCell data is publicly available at https://opencell.sf.czbiohub.org/download.

## A Technical Appendix

### A.1 Initialization

Models are initialized from LlamaGen [17] checkpoints pretrained on ImageNet class-conditional generation. New learnable embeddings for SOC, EOC, and EOS tokens are initialized from *N* (0, 0.02), and the conditioning embedder is randomly initialized.

### A.2 Classifier-free guidance

During training, with probability *p*_uncond_ = 0.1, we randomly replace the conditioning embedding with a learned unconditional embedding initialized as **c**_∅_ ∼ *N* (0, 1*/d*_ESM_). At inference, we interpolate between conditional and unconditional logits:

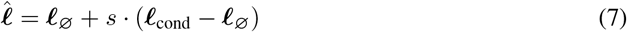

where *s* ≥ 1 is the guidance scale (*s* = 1 is equivalent to sampling without guidance).

### A.3 Inference

We condition on the first *C*_prefix_ channels and the ESM embedding of the protein sequence and autoregressively generate the remaining protein channel. Sampling uses top-*k* filtering with *k* = 2,000 and temperature *τ* = 1.0. Samples are generated without classifier-free guidance, unless otherwise specified. During inference, the transformer can flexibly generate new tokens given variable initial numbers of channels.

### A.4 Image Processing

For each FOV, every channel was loaded independently and resized from their native resolution (2048×2048) to 256×256 via Lanczos resampling. We additionally applied percentile-based intensity rescaling: pixel intensities were clipped to the 1st–99th percentile of the channel and divided by the per-channel maximum to yield float32 values in [0, 1].

### A.5 Training Strategy

All models are trained for 150 epochs with a batch size of 96 and the AdamW [29] optimizer. A peak learning rate of 1e-4 is used with *β*_1_ = 0.9, *β*_2_ = 0.95, and weight decay *λ* = 0.05. Learning rate follows linear warmup for the first 10 epochs, holding peak LR for 15% of the training budget, followed by linear decay. Gradient norms are clipped to 1.0. Training is performed with DistributedDataParallel (DDP) on a high-performance computing cluster. We use 1–2 A100 GPUs (Vermeer-B, Vermeer-L) or 6 H100 GPUs (Vermeer-XL), with 4 CPU workers per GPU and 10 GB RAM allocated per worker.

### A.6 Data Augmentations

For each multi-channel microscopy image, we pre-extract VQ-VAE codes under a fixed bank of deterministic augmentations and randomly sample one augmentation per epoch for training efficiency. Given an input of shape [H, W, 3, C] (C channels, e.g., nucleus, microtubule, ER, protein), we first upscale the center crop region to 1.1× the target resolution (282×282 for a 256×256 target), then apply torchvision’s TenCrop, which extracts the four corner crops and the center crop at the target resolution. We additionally apply horizontal flipping and random rotations of 0°, 90°, 180°, 270°, producing up to forty augmented views per image. The same crop index, flip, and rotation are applied consistently across all channels of a given image, preserving spatial registration across channels. Each view is independently tokenized by the frozen VQ-VAE and stored on disk. During training, one augmentation is randomly selected per sample.

### A.7 Dataset splitting

We constructed three disjoint evaluation regimes designed to isolate specific generalization axes. All splits were generated with a fixed random seed and grouped such that the two FOVs belonging to the same (cell line × protein) pair always remained in the same split.

- **Cell-line holdouts** All FOVs from three cell lines — GAMG, SK-MEL-30, and Hep-G2 — were removed from the pool before any other splitting and reserved as an out-of-distribution cell-line evaluation set. These cell-lines are chosen to test increasing difficulties of generalization to unseen cell-lines. GAMG is chosen as a cell-line from the same organ (brain) and cell-type (glioblastoma) as cell-lines in the training data. SK-MEL-30 is from the same organ (skin) but a different cell-type (melanoma) as cell-lines in the training data (carcinoma and keratinocyte are in the train set). Hep-G2 is chosen as a cell-line from an organ (liver) that’s not present in the training data
- **Unseen protein split** From the remaining pool, proteins were partitioned 90/10 at the protein level so that all FOVs of a given UniProt ID are assigned to exactly one split. The 10% holdout forms the unseen protein split, which contains proteins never observed during training. This partition is stratified on protein-level characteristics: primary localization for single-localizing proteins, and consistency type for multi-localizing proteins.
- **Unseen protein** × **cell-line split** From the 90% protein-train pool, we then performed a second 90/10 split, this time at the (protein × cell-line) group level, stratified jointly by cell-line and by protein-level characteristics (primary localization for single-localizing proteins or consistency type for multi-localizing proteins). Proteins may overlap between the final train set and this split, but the specific (protein × cell-line) combinations in this split are unseen during training. Groups whose stratification key appeared fewer than twice were split without stratification.

### A.8 Evaluation Metrics

We evaluate generated protein channels along three orthogonal axes: (i) distributional realism (FID), (ii) pixel- and intensity-level fidelity against the paired ground truth (MSE, intra-nuclear proportion with Pearson/Spearman), and (iii) semantic fidelity of the recovered subcellular localization pattern (a battery of multi-label classification metrics computed with a frozen SubCell classifier). All metrics are computed on the *protein channel only*, since the reference (nucleus / microtubule / ER) channels are copied from ground truth when generating in conditional mode and are therefore trivially matched.

#### A.8.1 Distributional realism

##### Fréchet Inception Distance (FID)

We compute FID with `torchmetrics.image.fid.FrechetInceptionDistance`. FID is the Fréchet distance between multivariate Gaussians fit to the pool-3 layer activations of an InceptionV3 model for both real and generated images:

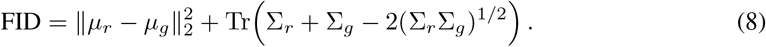

(*µ*_*r*_, Σ_*r*_) and (*µ*_*g*_, Σ_*g*_) are the mean and covariance of the features for the real and generated images respectively. FID captures whether the *set* of generated protein images is distributionally close to the real set, independent of per-sample pairing,.

#### A.8.2 Pixel- and intensity-level fidelity

##### Mean Squared Error (MSE)

Per-image MSE between paired real and generated protein images after rescaling and normalization to [0, 1]:

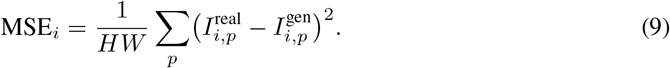

where *I* is pixel intensity, *p* indexes spatial pixels, and *H, W* are the height and width (in pixels) of the image. We report the mean MSE_*i*_ across matched pairs *i* of real/generated images. MSE penalizes coarse intensity- and spatial-pattern mismatch; however, it is sensitive to small spatial offsets and should be interpreted alongside the higher-level biological metrics below.

##### Intra-nuclear protein proportion (PCC intra-nuc prop)

For each image *i* we compute the fraction of the total protein signal that lies within the nucleus:

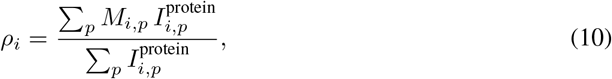

where *M*_*i*_ is a binary nuclear mask obtained by running StarDist 2D (`2D_versatile_fluo`, Gaussian smoothing with *σ* = 1.0) on the *true* nucleus channel. 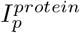 is the intensity at pixel *p* for the protein channel. The same mask is applied to both true and generated protein images.

We then report the Pearson correlation coefficient (PCC) over the paired vectors 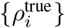 and 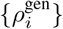. High PCC on this statistic indicates that the generator is not just producing plausible-looking protein textures, but actually redistributes signal between nuclear and cytoplasmic compartments in the correct direction for each conditioning input.

##### Localization fidelity

We compute metrics that assess whether the localization class of the generated protein stain is semantically consistent with the true protein stain. We do this by feeding each sample’s generated protein channel (paired with the true nucleus and microtubule channels) through **SubCellPortable**, a frozen multi-label localization classifier that outputs per-image probabilities over 31 subcellular compartment classes. We run SubCellPortable on both the true and generated images and compare its predictions against the human annotations provided by the Human Protein Atlas *y* ∈ {0, 1} ^31^, and — separately — compare classifier probabilities, on generated vs. true images to each other. We denote *s*_*i*_ ∈ [0, 1]^31^ as the SubCell probability vector on the generated image, and *ŝ*_*i*_ as the probability vector on the true image.

##### Top-1 accuracy

Fraction of samples for which the single highest-probability class output by SubCellPortable is among the HPA-annotated classes for that sample. Let *s*_*i,c*_ denote the predicted probability SubCellPortable to class c for sample *i* and *y*_*i*_ ∈ {0, 1} ^31^, the human ground-truth annotations for that sample:

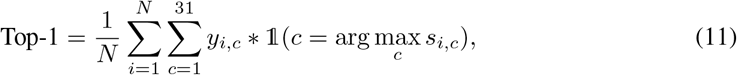

where 1(·) is the indicator function.

##### Top-3 accuracy

Same as above but with the top-3 predicted classes, counting a sample as correct if *any* of them overlaps with the HPA annotation.

##### Multi-label Ranking Average Precision (MLRAP)

Computed via `sklearn.metrics.label_ranking_average_precision_score`. For each sample, it averages, over the true positive labels, the precision among classes ranked at least as high as that label:

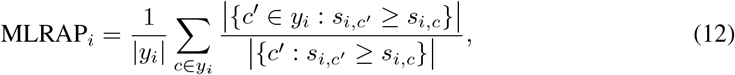

and reports the mean over samples. MLRAP ∈ (0, 1] with 1 being perfect; it measures whether the *true* localization classes are close to the top of the ranked probability list.

##### Coverage Error

Computed via `sklearn.metrics.coverage_error`: the average number of top-ranked classes one must traverse (from SubCell’s sorted probabilities) before all true positive labels have been retrieved. The minimum coverage error is the mean number of true labels per sample.

##### Macro / Micro Average Precision (AP)

Class-averaged area under the precision–recall (PR) curve from `sklearn.metrics.average_precision_score:`

- **Macro AP** averages per-class AP with equal weight, so rare localization classes contribute as much as frequent ones.
- **Micro AP** pools all (*y*_*i,c*_, *s*_*i,c*_) pairs across classes and samples into a single PR curve, giving weight proportional to class frequency.

##### Spearman correlation between true- and generated-image classifiers

For each sample, we compute the Spearman rank correlation between the full 31-dimensional probability vectors produced by SubCell on the true image (*ŝ*_*i*_) and on the generated image (*s*_*i*_):

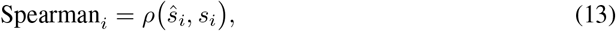

and report the mean over samples. Unlike the HPA-based metrics, this one does not require a ground-truth annotation and captures similarity of the full predicted localization profile, including patterns not annotated in HPA.

All of the above classifier-based metrics are reported twice (except for Spearman correlation): once with *s*_*i*_ taken from the generated image and once from the true image (as an upper-bound baseline that reflects the irreducible gap between the frozen classifier and the HPA annotations).

### A.9 Benchmarking Protocol

For Table 1 and Table 2, CELL-E 2 and CELL-Diff are run using publicly available checkpoints and following the authors’ instructions for preprocessing the inputs. Because of a reported issue in PUPS, where using composite JPEG inputs introduces cross-channel signal [8], we retrain the PUPS method, following the authors’ protocol, but instead using TIFF inputs and our own preprocessing strategy.

To normalize brightness intensity, which differs across methods due to preprocessing differences, the generated image and true image for each method are percentile-clipped at the 1st and 99th percentile and rescaled to [0,1]. This transformation is applied to the generated output and the true output as a post-processing step for all methods, including Vermeer.

We additionally retrain CELL-E 2 and CELL-Diff, following the authors’ protocol, on our own preprocessed data, which is in Table S5 and Table S6. For systematic comparison, we also retrain Vermeer at a image resolution matched to CELL-Diff, Vermeer-0.25-L (Table S5, S6).

### A.10 Protein Annotation

Using the Human Protein Atlas’ per-Field-of-view (FOV) localization annotations (mapped onto 19 canonical subcellular compartment classes), every protein was assigned (i) as single-localizing vs. multi-localizing — based on whether any FOV carried more than one localization label, and (ii) a consistency type capturing how its localization varies across imaging contexts:

- Invariant: all cell-lines and fields-of-view in the HPA exhibit the same localization.
- Deterministically variable: localization varies across cell lines but is consistent across FOVs within a given cell line.
- Stochastically variable: localization varies across fields-of-view within the same cell line.

## B Broader Impacts Statement

Vermeer is a generative image model trained on the Human Protein Atlas, which contains images of cancer cell-lines. While our work is methodological, there are potential societal impacts. Vermeer may support biological research by enabling the generation of realistic protein localization images, which could assist in hypothesis generation and prioritization of downstream experiments. However, the model may reflect biases present in the Human Protein Atlas dataset which could lead to systematic errors when the model is applied to underrepresented biological contexts. As a generative image model, Vermeer could potentially be misused to generate synthetic microscopy images that are difficult to distinguish from real data. This has potential risks, such as the fabrication of data or manipulation of experimental results. We require users to adhere to safe and responsible usage of Vermeer in our model license.

## C Supplemental Tables

**Table S1:**
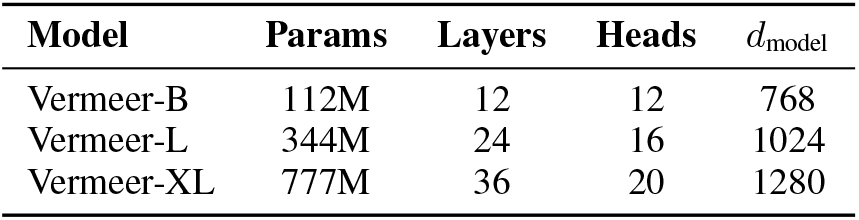
Vermeer model configurations.

| **Model** | **Params** | **Layers** | **Heads** | *d*<sub>model</sub> |
|---|---|---|---|---|
| Vermeer-B | 112M | 12 | 12 | 768 |
| Vermeer-L | 344M | 24 | 16 | 1024 |
| Vermeer-XL | 777M | 36 | 20 | 1280 |

**Table S2:** SubCell oracle image classifier metrics on unseen protein × cell-line split, scored against HPA labels. Baselines are run with Vermeer-XL. AP: attention pooled, CA: channel augmented.

| Model | MLRAP<br>(↑) | Coverage error<br>(↓) | Macro AP<br>(↑) | Micro AP<br>(↑) | Top 3 Accuracy<br>(↑) | Top 1 Accuracy<br>(↑) |
|---|---|---|---|---|---|---|
| Vermeer-B | 0.621 | 5.96 | 0.150 | 0.464 | 0.761 | 0.579 |
| Vermeer-L | 0.660 | 5.34 | 0.188 | 0.520 | 0.792 | 0.627 |
| Vermeer-L AP | 0.659 | 5.34 | **0.192** | 0.525 | 0.795 | 0.625 |
| Vermeer-XL | **0.662** | **5.28** | 0.190 | **0.526** | **0.796** | **0.629** |
| Vermeer-XL CA | 0.635 | 5.60 | 0.167 | 0.487 | 0.775 | 0.590 |
| Pixel shuffled (baseline) | 0.380 | 8.47 | 0.0653 | 0.203 | 0.659 | 0.153 |
| Protein prefix shuffled (baseline) | 0.460 | 8.00 | 0.0591 | 0.254 | 0.630 | 0.351 |
| True | 0.693 | 4.86 | 0.256 | 0.578 | 0.813 | 0.664 |

**Table S3:**
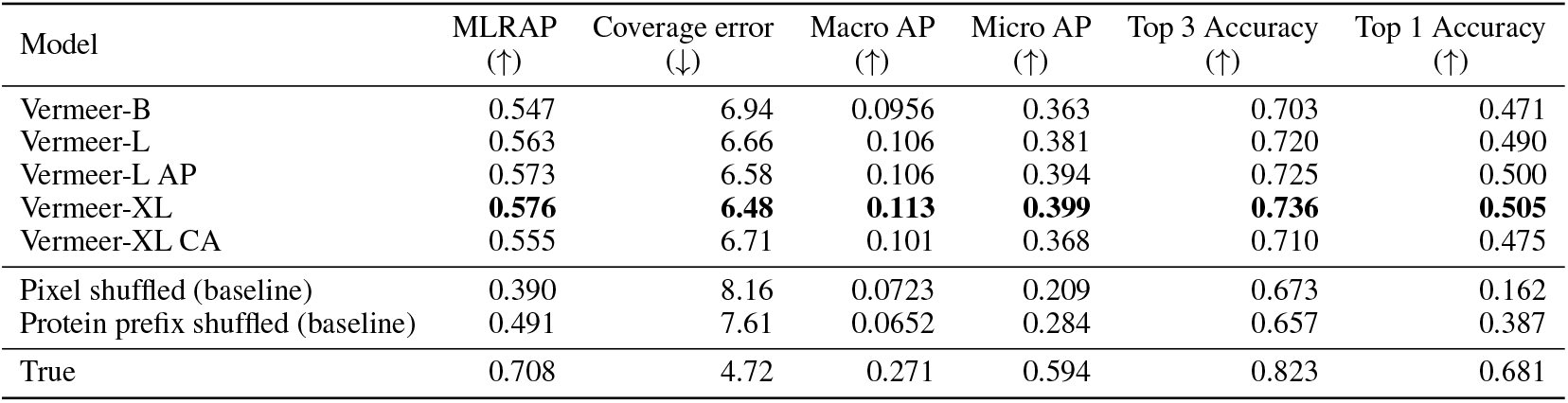
SubCell oracle image classifier metrics on unseen protein split, scored against HPA labels. Baselines are run with Vermeer-XL. AP: attention pooled, CA: channel augmented.

**Table S4:** SubCell oracle image classifier metrics on cell-line holdouts split, scored against HPA labels. Baselines are run with Vermeer-XL. AP: attention pooled, CA: channel augmented.

| Model | MLRAP<br>(↑) | Coverage error<br>(↓) | Macro AP<br>(↑) | Micro AP<br>(↑) | Top 3 Accuracy<br>(↑) | Top 1 Accuracy<br>(↑) |
|---|---|---|---|---|---|---|
| Vermeer-B | 0.570 | 6.84 | 0.144 | 0.412 | 0.712 | 0.510 |
| Vermeer-L | **0.616** | 6.18 | 0.197 | **0.478** | 0.744 | **0.578** |
| Vermeer-L AP | 0.615 | **6.09** | 0.188 | 0.475 | **0.750** | 0.571 |
| Vermeer-XL | 0.613 | 6.11 | **0.199** | 0.473 | 0.738 | 0.573 |
| Vermeer-XL CA | 0.575 | 6.54 | 0.164 | 0.424 | 0.710 | 0.513 |
| Pixel shuffled (baseline) | 0.366 | 8.82 | 0.0798 | 0.201 | 0.615 | 0.165 |
| Protein prefix shuffled (baseline) | 0.463 | 8.14 | 0.0703 | 0.264 | 0.622 | 0.360 |
| True | 0.673 | 5.06 | 0.255 | 0.562 | 0.794 | 0.639 |

**Table S5:**
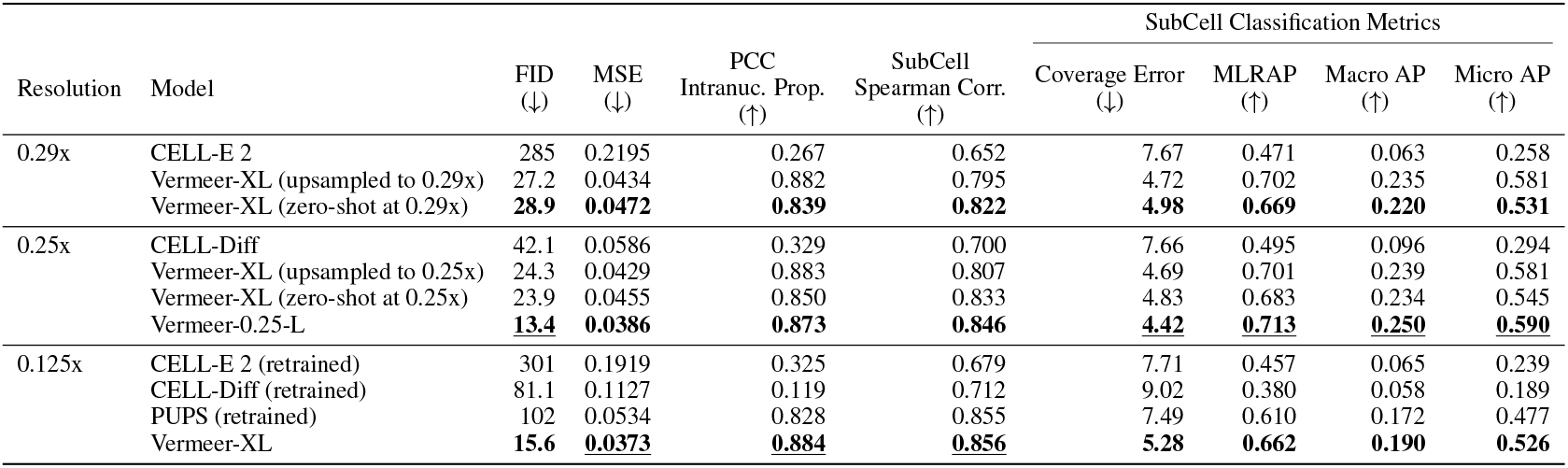
Benchmarking results on unseen protein × cell line split, grouped by resolution. Upsampled indicates upsampling both the true and generated Vermeer images to the target resolution. Zero-shot indicates running Vermeer virtual staining directly at the target resolution. Vermeer-0.25-L indicates Vermeer retrained at 0.25x. Retrained indicates baselines retrained on our data split and preprocessing at 0.125x. Best metrics within each resolution block are bolded (excluding upsampled); overall best are underlined.

| Resolution | Model | FID<br>(↓) | MSE<br>(↓) | PCC<br>Intranuc. Prop.<br>(↑) | SubCell<br>Spearman Corr.<br>(↑) | SubCell Classification Metrics |  |  |  |
|---|---|---|---|---|---|---|---|---|---|
|  |  |  |  |  |  | Coverage Error<br>(↓) | MLRAP<br>(↑) | Macro AP<br>(↑) | Micro AP<br>(↑) |
| 0.29x | CELL-E 2 | 285 | 0.2195 | 0.267 | 0.652 | 7.67 | 0.471 | 0.063 | 0.258 |
|  | Vermeer-XL (upsampled to 0.29x) | 27.2 | 0.0434 | 0.882 | 0.795 | 4.72 | 0.702 | 0.235 | 0.581 |
|  | Vermeer-XL (zero-shot at 0.29x) | **28.9** | **0.0472** | **0.839** | **0.822** | **4.98** | **0.669** | **0.220** | **0.531** |
| 0.25x | CELL-Diff | 42.1 | 0.0586 | 0.329 | 0.700 | 7.66 | 0.495 | 0.096 | 0.294 |
|  | Vermeer-XL (upsampled to 0.25x) | 24.3 | 0.0429 | 0.883 | 0.807 | 4.69 | 0.701 | 0.239 | 0.581 |
|  | Vermeer-XL (zero-shot at 0.25x) | 23.9 | 0.0455 | 0.850 | 0.833 | 4.83 | 0.683 | 0.234 | 0.545 |
|  | Vermeer-0.25-L | 13.4 | **0.0386** | **0.873** | **0.846** | 4.42 | 0.713 | 0.250 | 0.590 |
| 0.125x | CELL-E 2 (retrained) | 301 | 0.1919 | 0.325 | 0.679 | 7.71 | 0.457 | 0.065 | 0.239 |
|  | CELL-Diff (retrained) | 81.1 | 0.1127 | 0.119 | 0.712 | 9.02 | 0.380 | 0.058 | 0.189 |
|  | PUPS (retrained) | 102 | 0.0534 | 0.828 | 0.855 | 7.49 | 0.610 | 0.172 | 0.477 |
|  | Vermeer-XL | **15.6** | 0.0373 | 0.884 | 0.856 | **5.28** | **0.662** | **0.190** | **0.526** |

**Table S6:** Benchmarking results on unseen protein split, grouped by resolution. Upsampled indicates upsampling both the true and generated Vermeer images to the target resolution. Zero-shot indicates running Vermeer virtual staining directly at the target resolution. Vermeer-0.25-L indicates Vermeer retrained at 0.25x. Retrained indicates baselines retrained on our data split and preprocessing at 0.125x. Best metrics within each resolution block are bolded (excluding upsampled); overall best are underlined.

| Resolution | Model | FID<br>(↓) | MSE<br>(↓) | PCC<br>Intranuc. Prop.<br>(↑) | SubCell<br>Spearman Corr.<br>(↑) | SubCell Classification Metrics |  |  |  |
|---|---|---|---|---|---|---|---|---|---|
|  |  |  |  |  |  | Coverage Error<br>(↓) | MLRAP<br>(↑) | Macro AP<br>(↑) | Micro AP<br>(↑) |
| 0.29x | CELL-E 2 | 285 | 0.2220 | 0.287 | 0.649 | 7.44 | 0.483 | 0.066 | 0.267 |
|  | Vermeer-XL (upsampled to 0.29x) | 28.5 | 0.0560 | 0.632 | 0.744 | 6.05 | 0.600 | 0.129 | 0.423 |
|  | Vermeer-XL (zero-shot at 0.29x) | **31.3** | **0.0566** | **0.589** | **0.770** | **6.25** | **0.581** | **0.124** | **0.398** |
| 0.25x | CELL-Diff | 41.9 | 0.0589 | 0.340 | 0.699 | 7.45 | 0.506 | 0.092 | 0.307 |
|  | Vermeer-XL (upsampled to 0.25x) | 25.4 | 0.0553 | 0.629 | 0.754 | 6.06 | 0.599 | 0.130 | 0.424 |
| 0.25x | CELL-Diff | 41.9 | 0.0589 | 0.340 | 0.699 | 7.45 | 0.506 | 0.092 | 0.307 |
|  | Vermeer-XL (upsampled to 0.25x) | 25.4 | 0.0553 | 0.629 | 0.754 | 6.06 | 0.599 | 0.130 | 0.424 |
|  | Vermeer-XL (zero-shot at 0.25x) | 25.4 | 0.0554 | 0.598 | 0.776 | **<u>6.21</u>** | 0.587 | **<u>0.125</u>** | 0.406 |
|  | Vermeer-0.25-L | **<u>15.3</u>** | **0.0488** | **0.610** | **0.777** | 6.22 | **<u>0.590</u>** | 0.110 | **<u>0.408</u>** |
| 0.125x | CELL-E 2 (retrained) | 300 | 0.1924 | 0.307 | 0.677 | 7.52 | 0.466 | 0.066 | 0.241 |
|  | CELL-Diff (retrained) | 83.3 | 0.1150 | 0.119 | 0.706 | 8.64 | 0.388 | 0.062 | 0.194 |
|  | PUPS (retrained) | 104 | 0.0691 | **<u>0.652</u>** | **<u>0.825</u>** | 8.28 | 0.538 | 0.092 | 0.369 |
|  | Vermeer-XL | **18.1** | **<u>0.0470</u>** | 0.624 | 0.802 | **6.48** | **0.576** | **0.113** | **0.399** |

**Table S7:** Ablation of protein language model conditioning. Vermeer-L (no ESM) is trained with learnable protein embeddings. For the unseen protein split, a nearest neighbor search is run in ESM embedding space using cosine similarity, and the learnable embedding for the closest train protein is used. Best metrics within each split are bolded.

|  | Unseen Proteins |  |  | Unseen Cell Line × Protein |  |  |
|---|---|---|---|---|---|---|
| Model | MSE<br>(↓) | PCC<br>Intranuc. Prop.<br>(↑) | Oracle<br>Spearman Corr.<br>(↑) | MSE<br>(↓) | PCC<br>Intranuc. Prop.<br>(↑) | Oracle<br>Spearman Corr.<br>(↑) |
| Vermeer-L (no ESM) | 0.0539 | 0.460 | 0.768 | **0.0387** | **0.866** | **0.847** |
| Vermeer-L | **0.0514** | **0.580** | **0.790** | 0.0401 | 0.852 | 0.842 |

## Supplemental Figures

**Figure S1:**
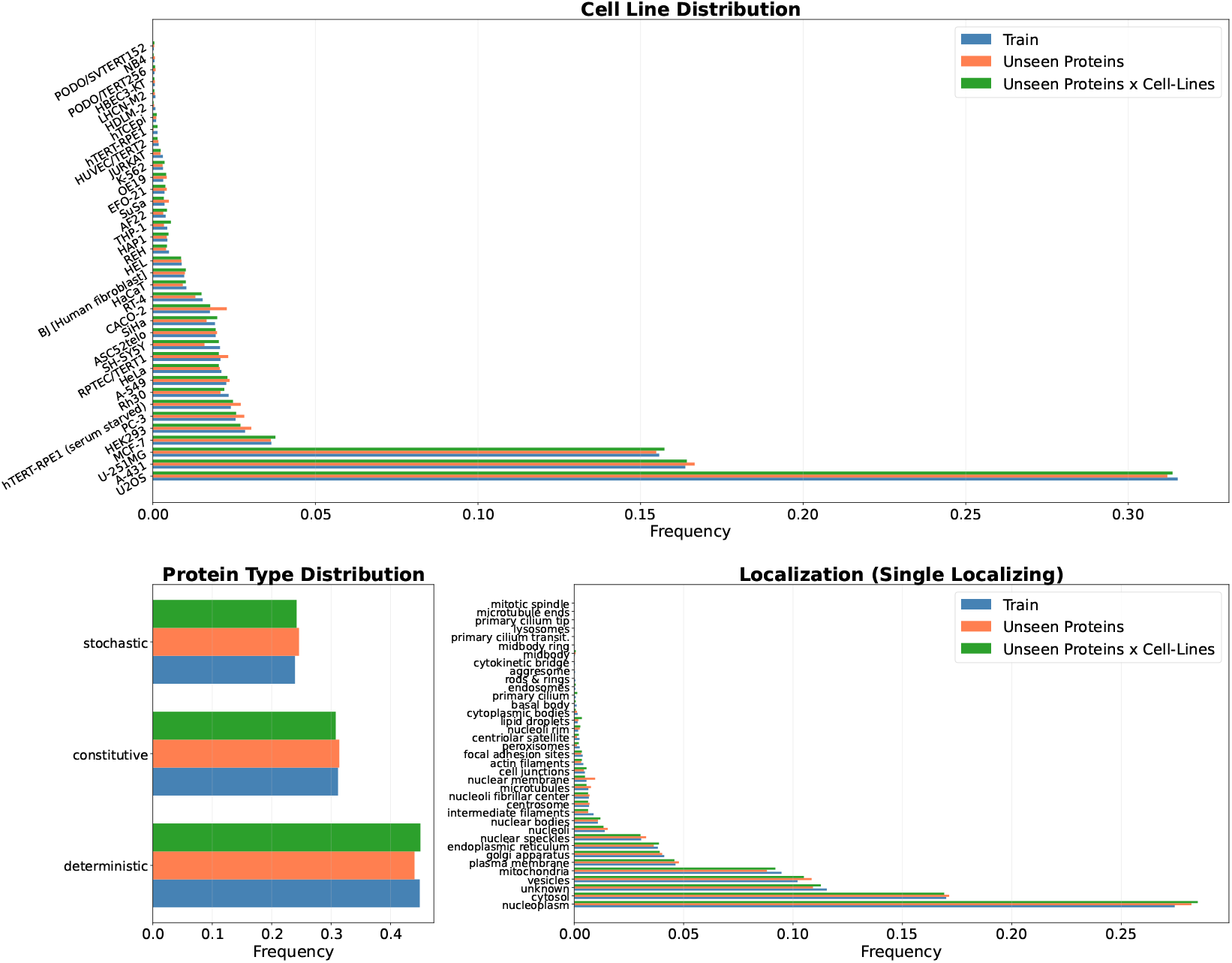
Cell-line and protein characteristic (variability and localization) distributions across train, unseen protein, and unseen protein × cell-line splits. Train: 74347 samples, 11661 proteins. Unseen protein split: 9314 samples, 1306 proteins. Unseen protein × cell-line split: 8275 samples, 3174 proteins.

**Figure S2:**
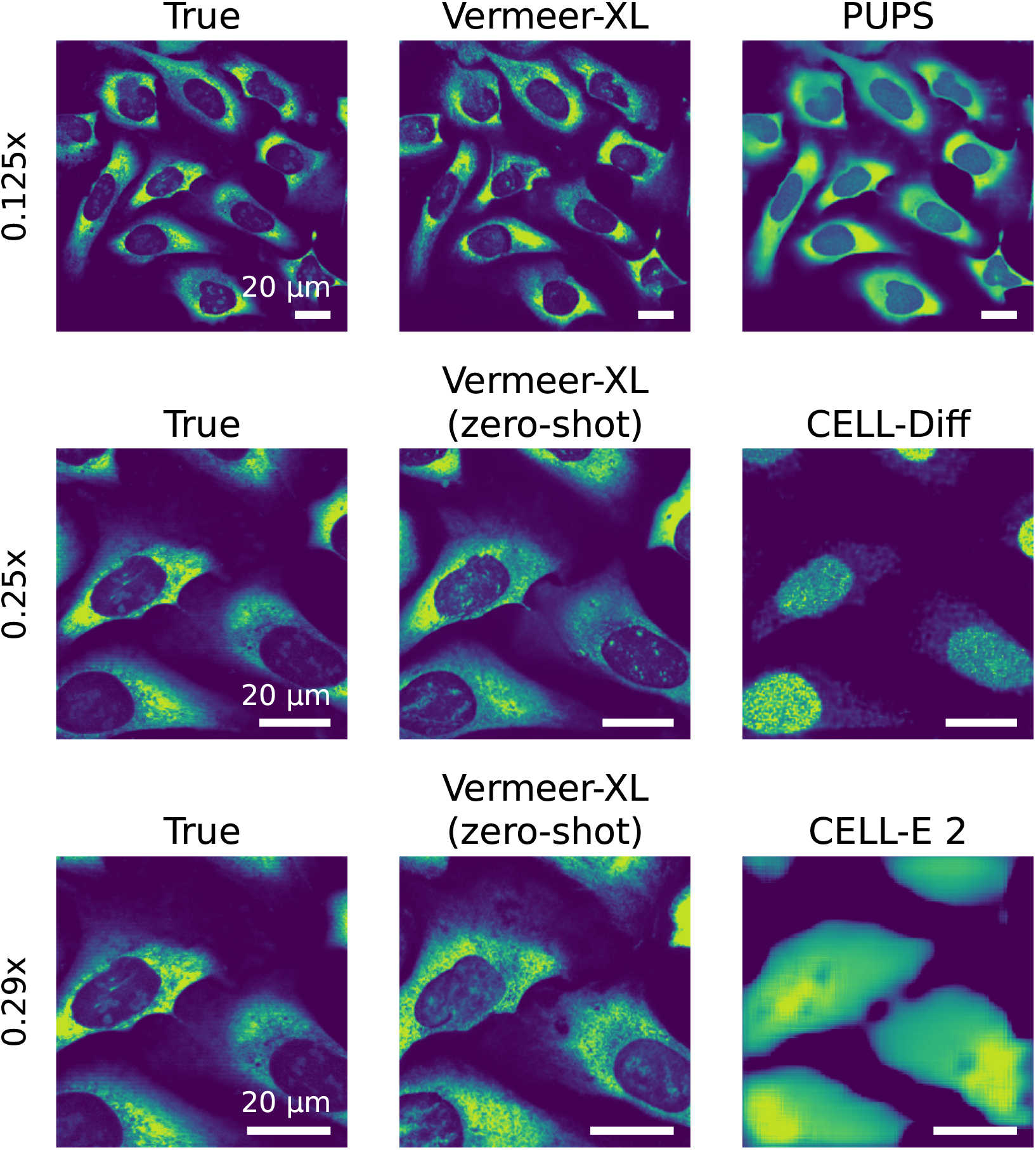
Example of protein virtual staining generations across different methods. Different methods operate at different resolutions, so we additionally include Vermeer zero-shot at various resolutions.

**Figure S3:**
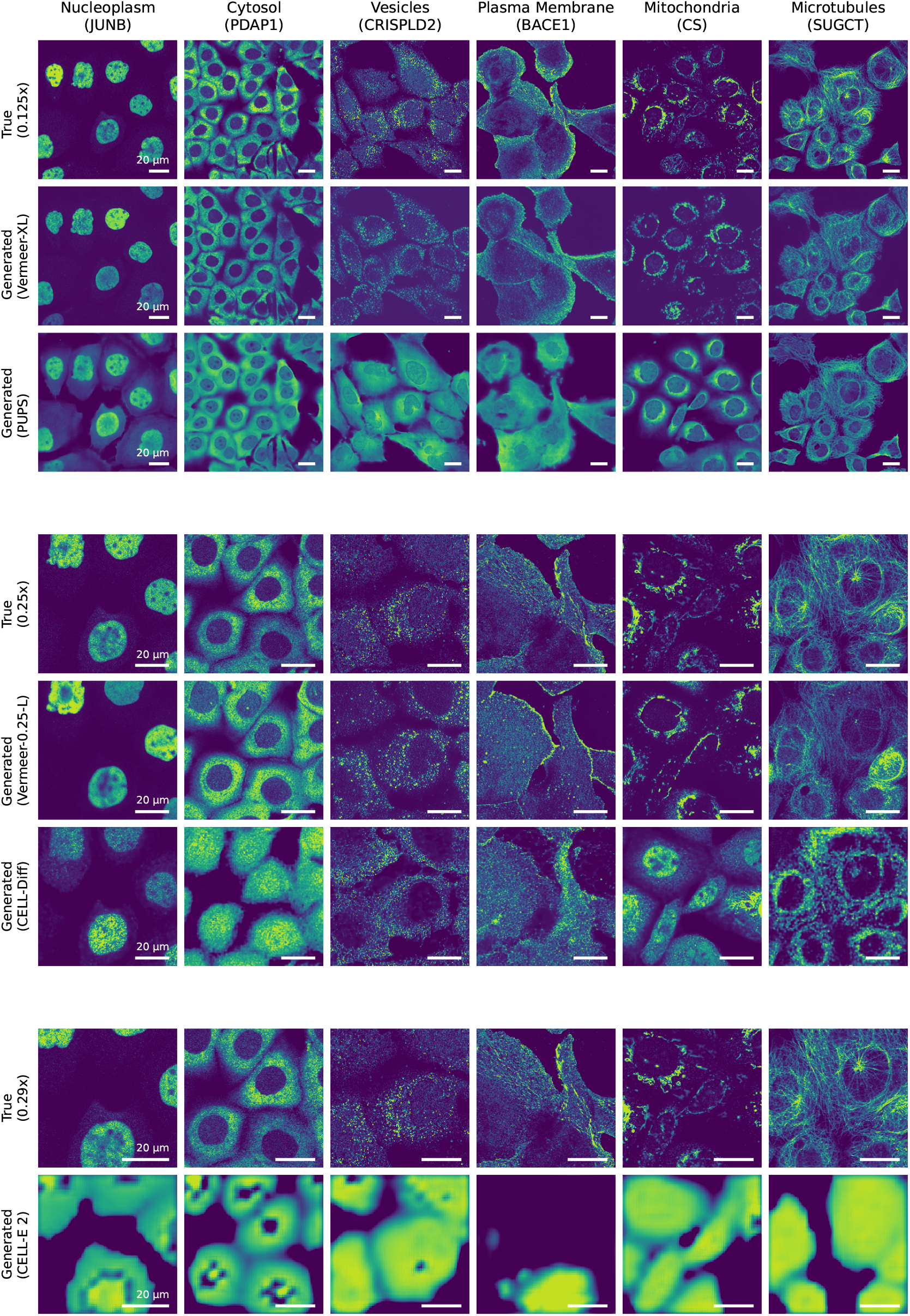
Generations across localization classes for Vermeer and existing methods on unseen protein x cell-line split.

**Figure S4:**
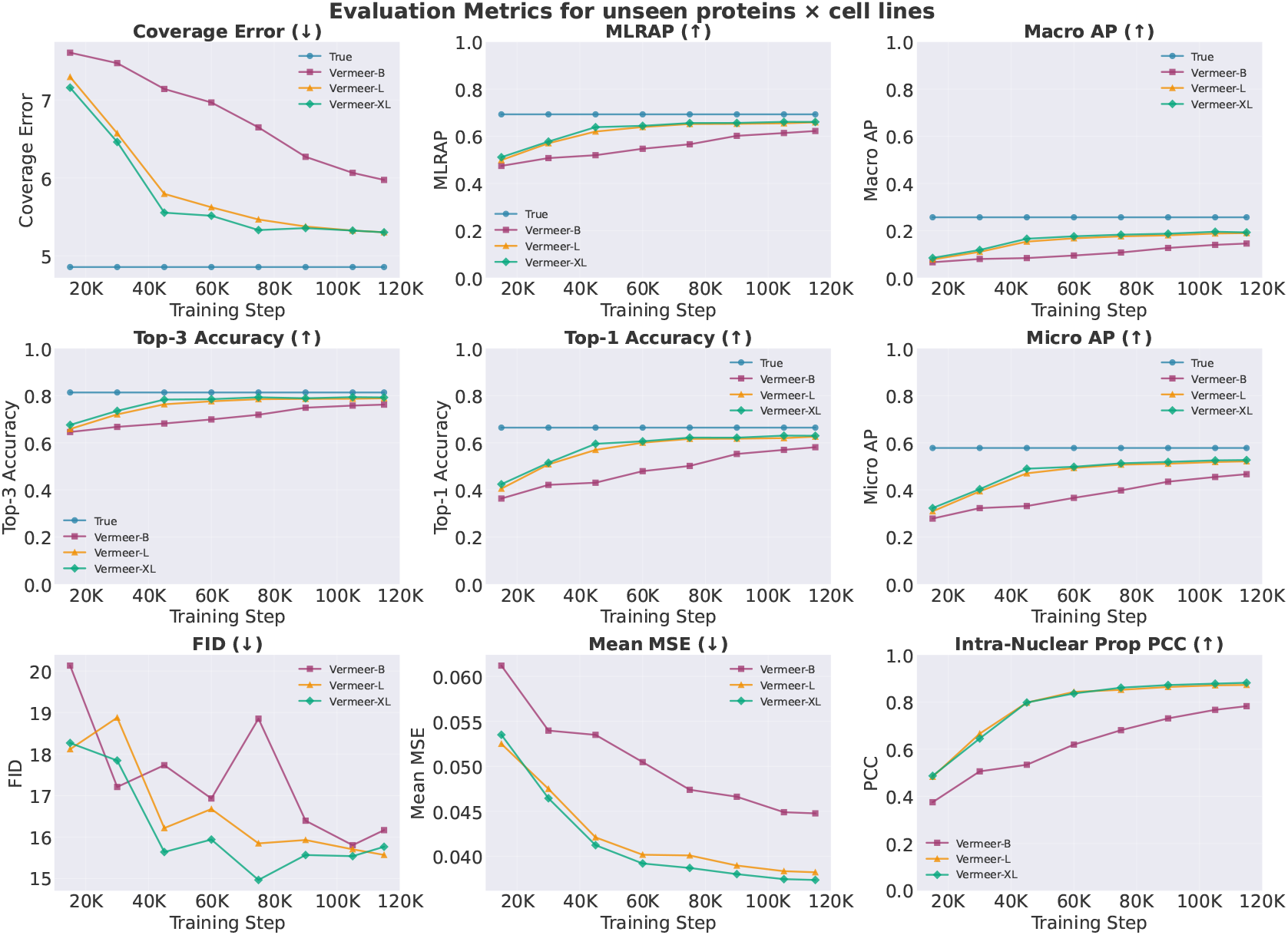
Evaluation metrics across training steps for Vermeer-B, Vermeer-L, and Vermeer-XL models on unseen protein × cell-line split

**Figure S5:**
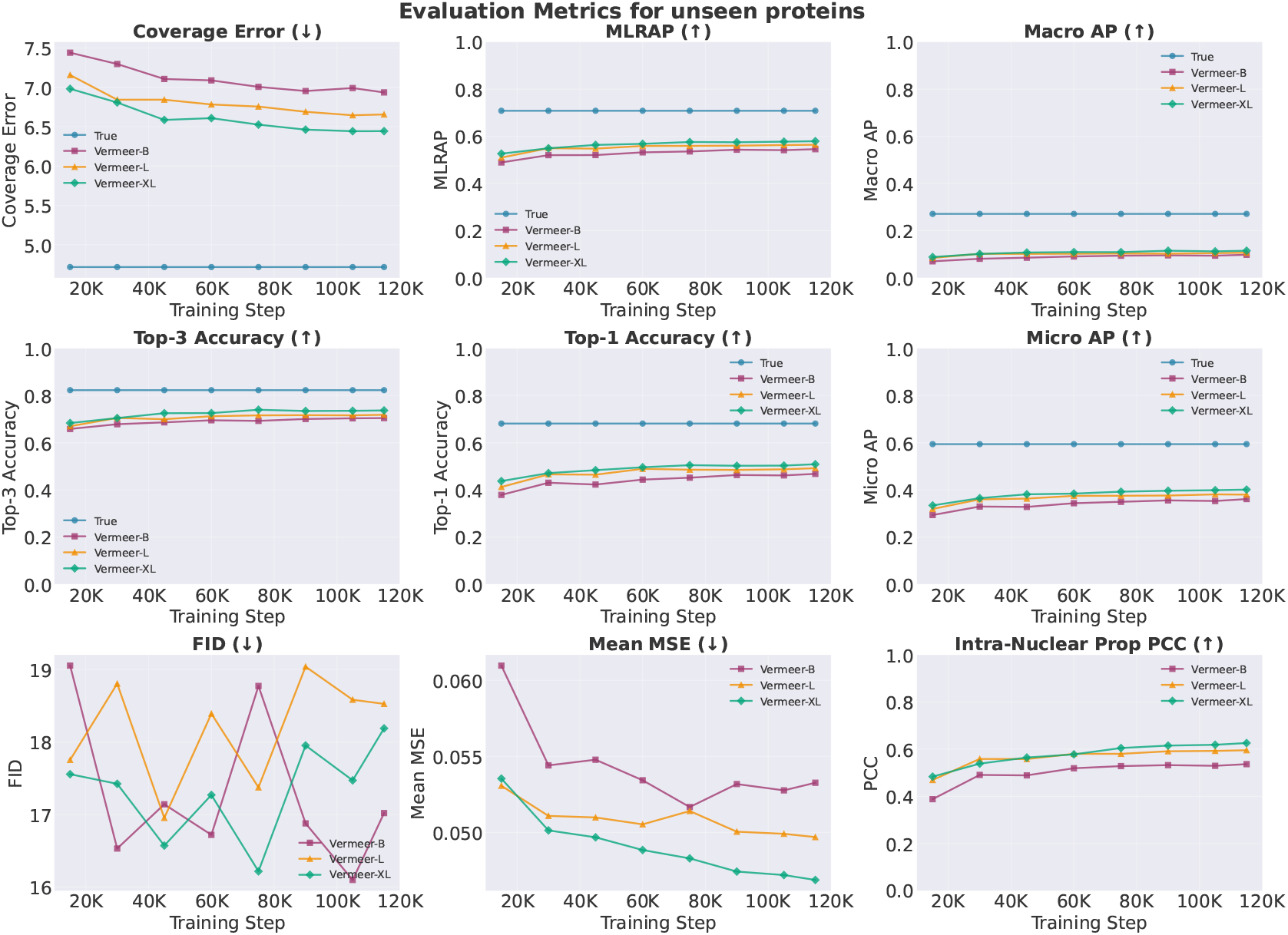
Evaluation metrics across training steps for Vermeer-B, Vermeer-L, and Vermeer-XL models on unseen protein split

**Figure S6:**
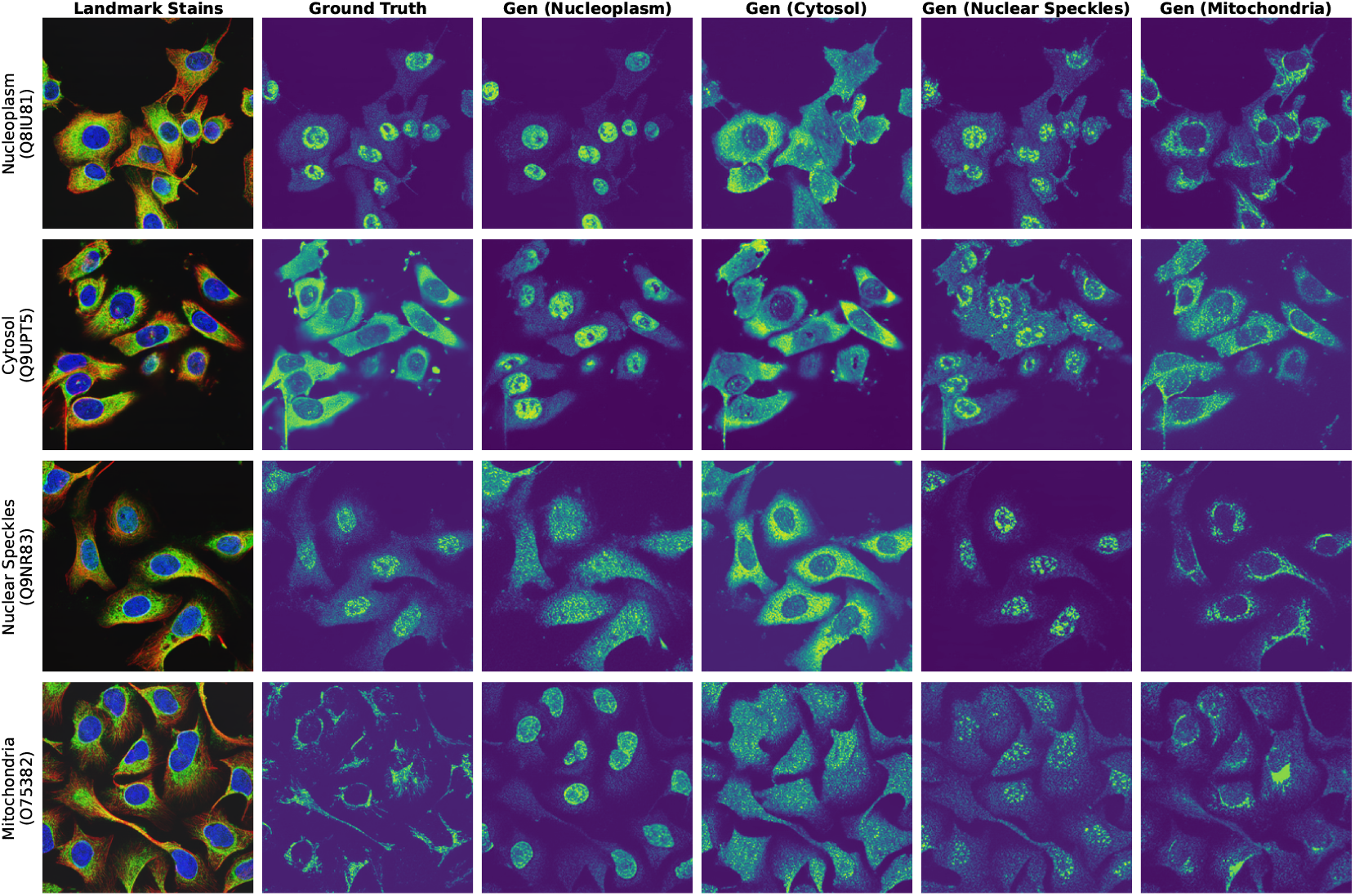
Confirmation that the model does not rely on spectral bleedthrough by generating multiple protein stains from the same landmark stains for different proteins. All combinations of proteins and landmark stains are shown.

**Figure S7:**
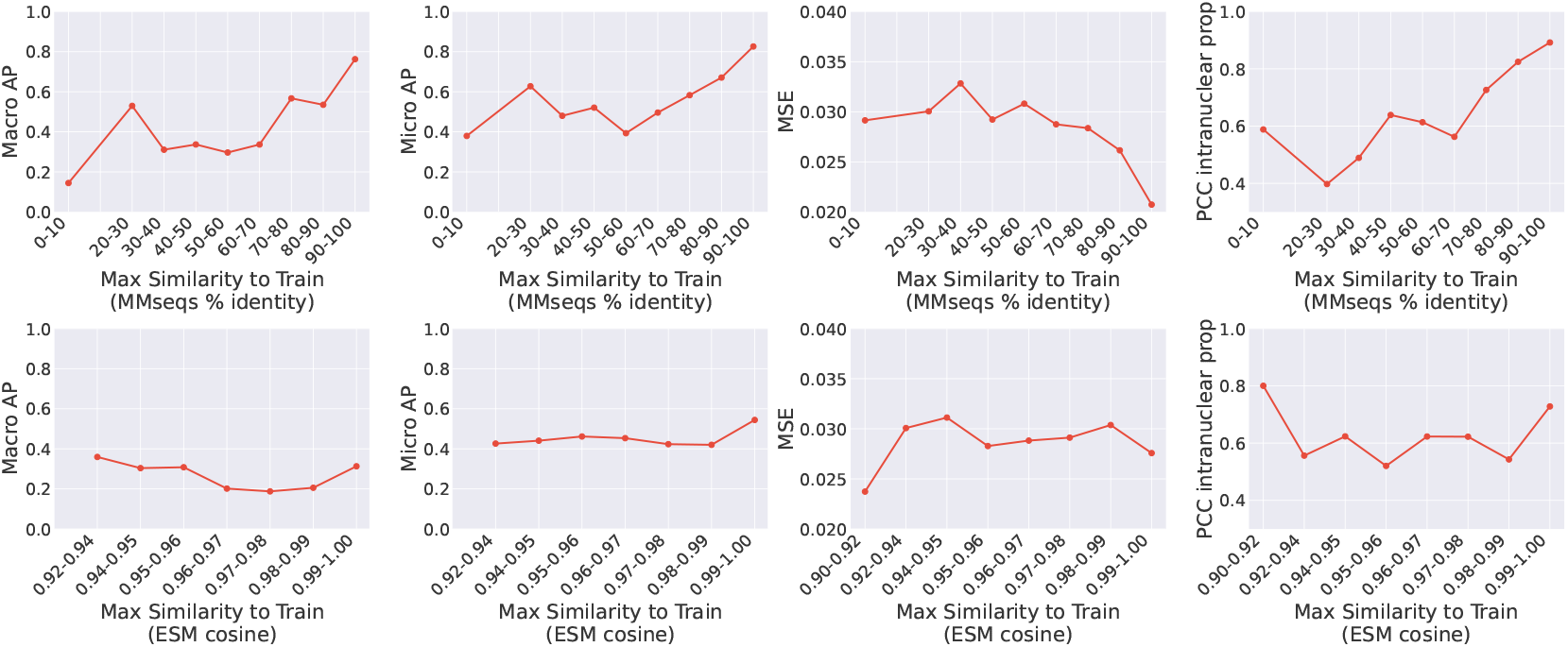
Evaluation metrics versus sequence similarity. Top panel displays sequence similarity calculated by MMseqs2 (defined as max similarity of validation sequence to any train sequence). Sequence similarity is binned into deciles. Bottom panel displays sequence similarity as the maximum cosine similarity between the ESM embedding of a given validation sequence and any train sequence. Cosine similarity is binned into two-percentile groups. Macro and Micro AP are computed after filtering to localization classes which have at least 50 examples in the validation set, to reduce noise in the trend-line.

**Figure S8:**
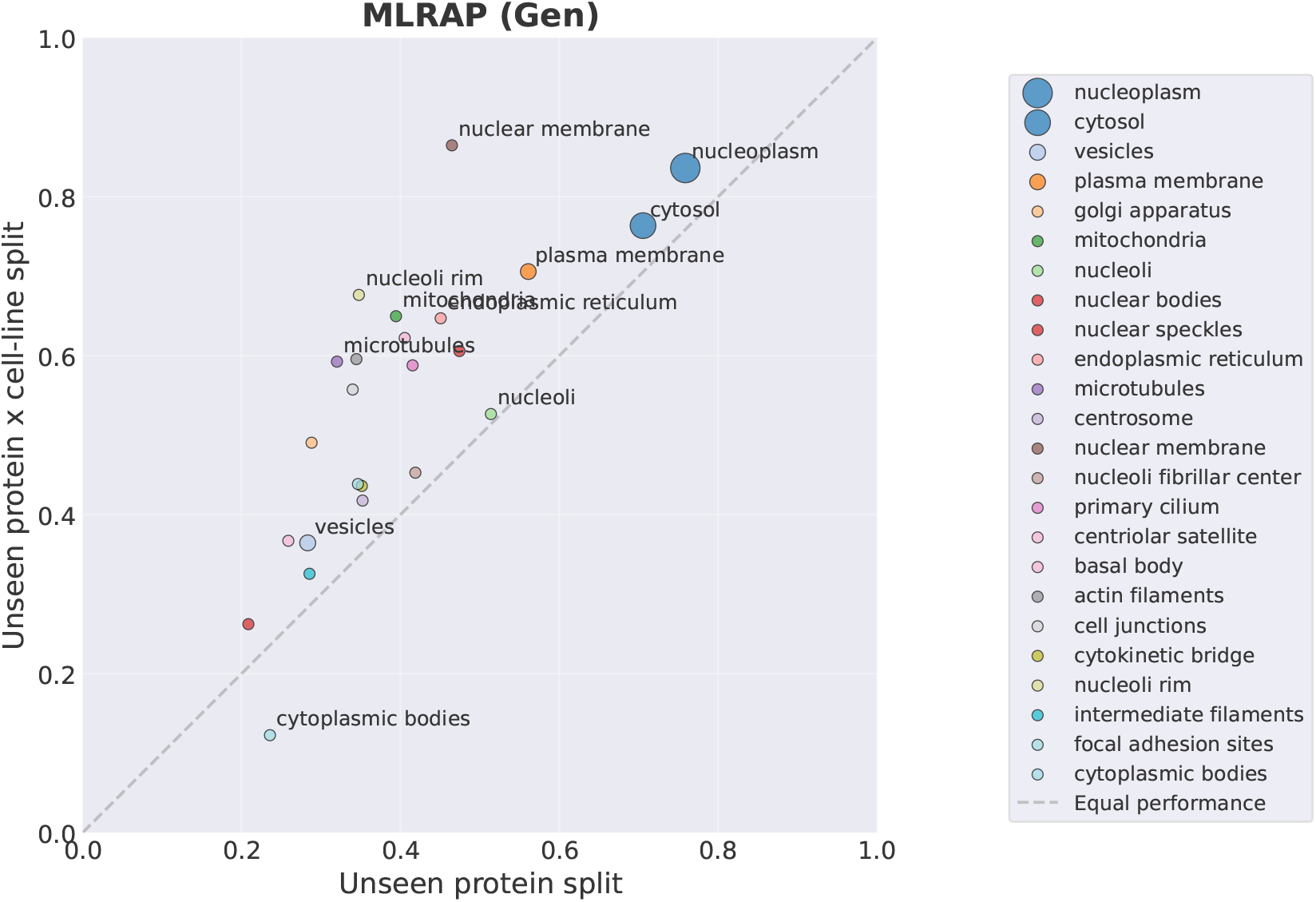
Scatter plot of evaluation metrics, stratified by localization class for unseen protein split (on x-axis) and unseen protein × cell-line split (on y-axis). Dot size indicates frequency of localization class.

**Figure S9:**
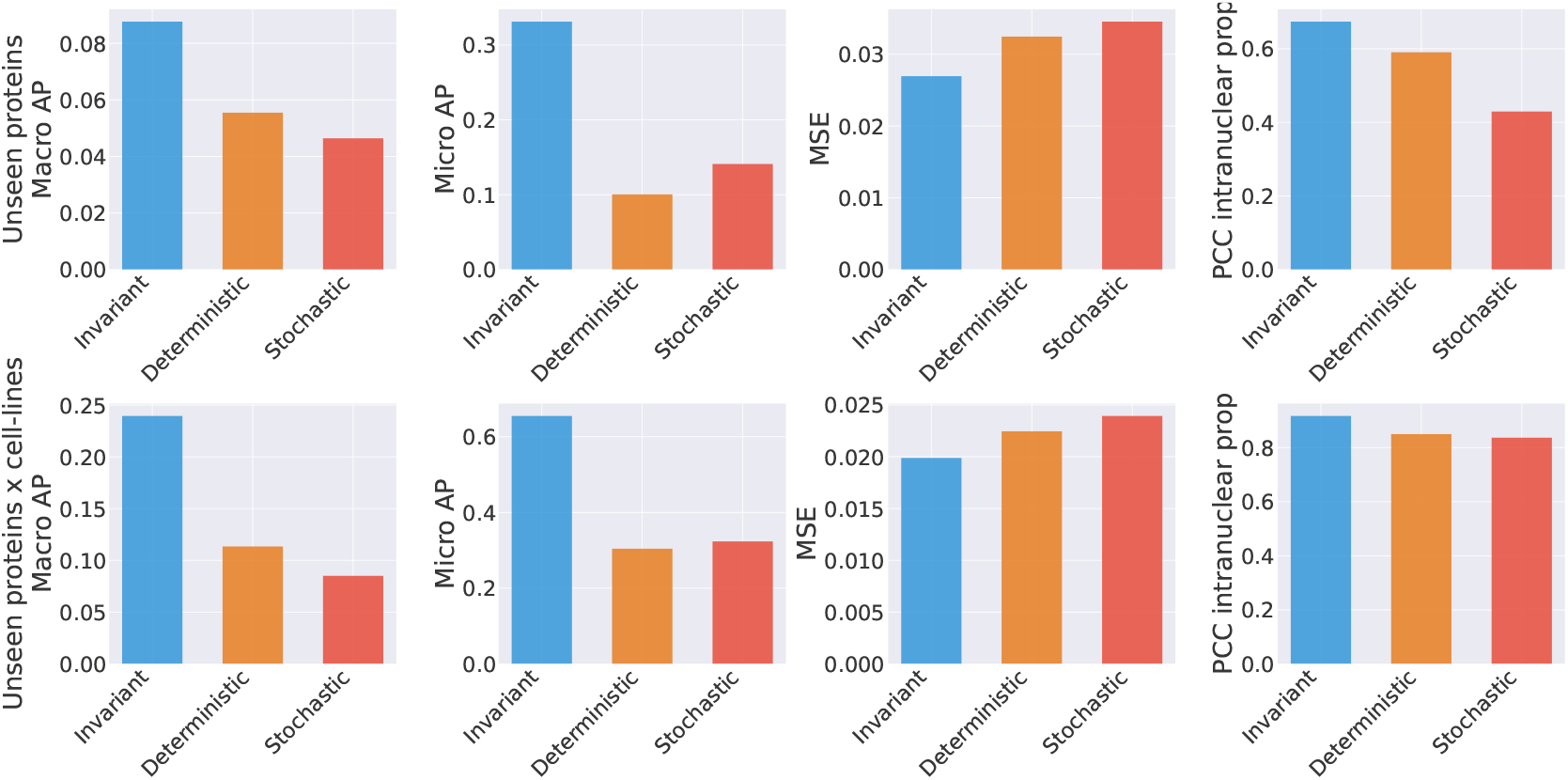
Evaluation metrics stratified by protein localization variability. Top panel shows unseen protein split and bottom panel shows unseen protein × cell-line split.

**Figure S10:**
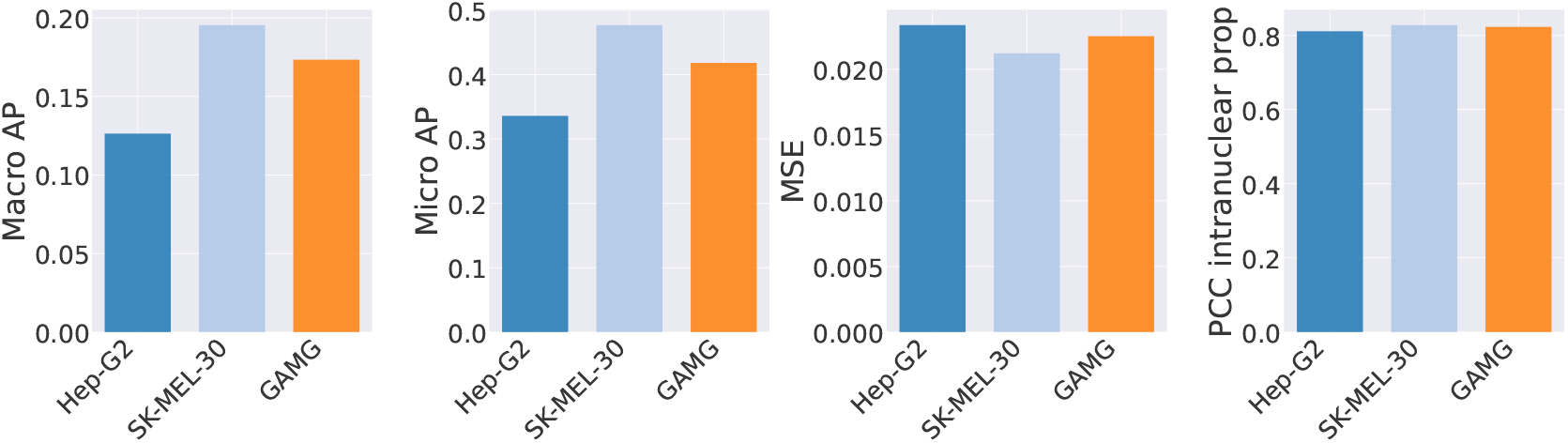
Evaluation metrics stratified by cell-line in unseen cell-line split.

**Figure S11:**
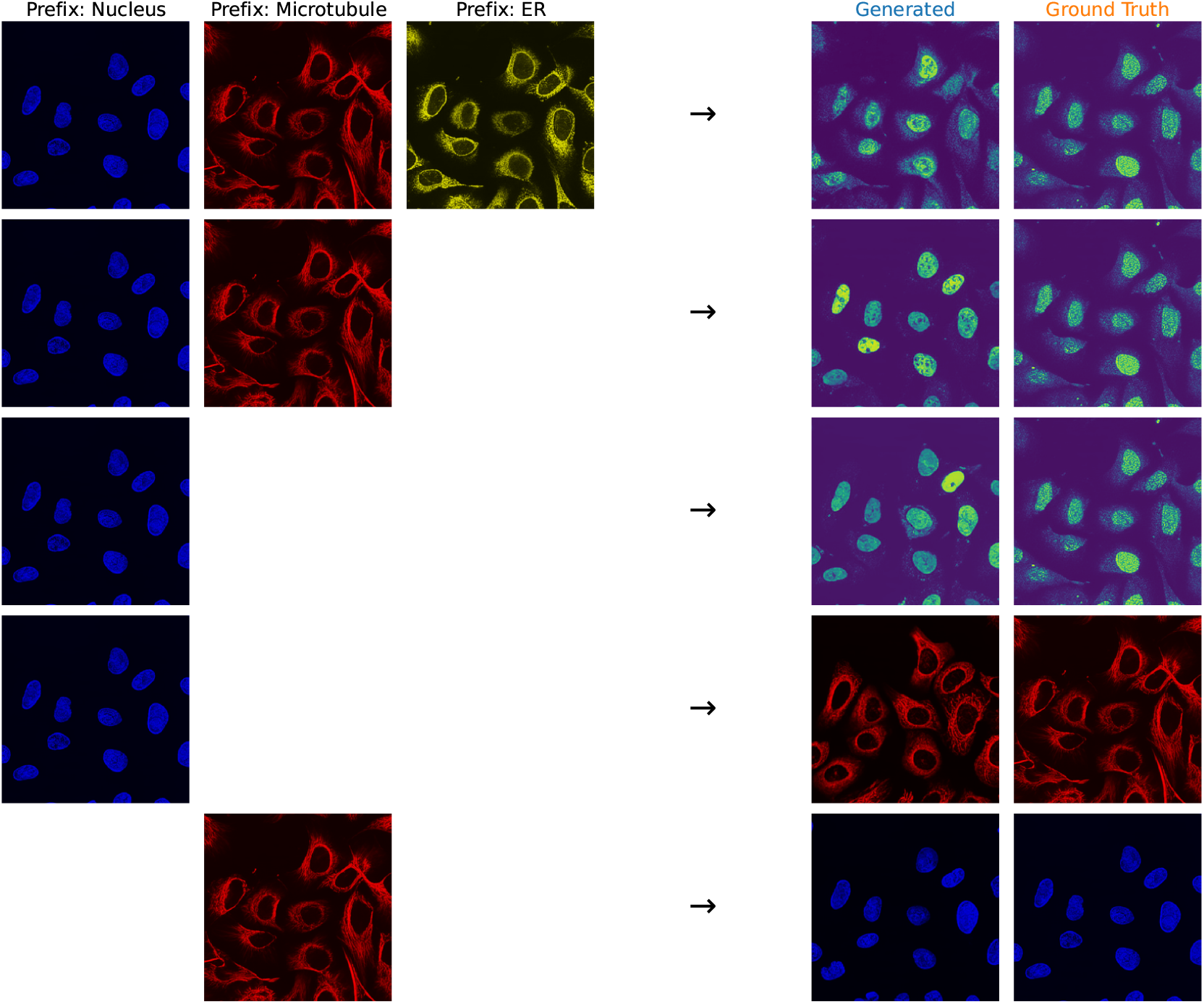
Demonstration of Vermeer’s channel adaptive generation capabilities. Left shows channels given as input and right shows generation, along with ground truth

## References

Alina Sigaeva, Charlotte Hutchings, Anthony Cesnik, Kathryn S. Lilley, and Emma Lundberg. Subcellular localization as a driver of protein function. Nature Reviews Molecular Cell Biology, 27(7):493–510, 2026. doi: 10.1038/s41580-026-00947-3.

Jessica Lacoste et al. Pervasive mislocalization of pathogenic coding variants underlying human disorders. Cell, 187(23):6725–6741.e13, 2024. doi: 10.1016/j.cell.2024.09.003.

Peter J. Thul et al. A subcellular map of the human proteome. Science, 356(6340):eaal3321, 2017. doi: 10.1126/science.aal3321.

Ankit Gupta et al. SubCell: Proteome-aware vision foundation models for microscopy capture single-cell biology. bioRxiv preprint, 2024. doi: 10.1101/2024.12.06.627299.

Judice L Y Koh et al. CYCLoPs: A Comprehensive Database Constructed from Automated Analysis of Protein Abundance and Subcellular Localization Patterns in Saccharomyces cerevisiae. G3 Genes|Genomes|Genetics, 5(6):1223–1232, 2015. doi: 10.1534/g3.115.017830.

Anastasia Razdaibiedina et al. PIFiA: Self-supervised approach for protein functional annotation from single-cell imaging data. Molecular Systems Biology, 20(5):521–548, 2024. doi: 10.1038/s44320-024-00029-6.

Uri Weill et al. Genome-wide SWAp-Tag yeast libraries for proteome exploration. Nature Methods, 15(8):617–622, 2018. doi: 10.1038/s41592-018-0044-9.

Xinyi Zhang, Yitong Tseo, Yunhao Bai, Fei Chen, and Caroline Uhler. Prediction of protein subcellular localization in single cells. Nature Methods, 22(6):1265–1275, 2025. doi: 10.1038/s41592-025-02696-1.

Emaad Khwaja, Yun Song, Aaron Agarunov, and Bo Huang. CELL-E 2: Translating Proteins to Pictures and Back with a Bidirectional Text-to-Image Transformer. In Advances in Neural Information Processing Systems 36, pages 4899–4914, New Orleans, Louisiana, USA, 2023. Neural Information Processing Systems Foundation, Inc. (NeurIPS). doi: 10.52202/075280-0217.

Dihan Zheng and Bo Huang. Bridging Protein Sequences and Microscopy Images with Unified Diffusion Models. In Proceedings of the 42nd International Conference on Machine Learning, pages 78139–78155. PMLR, 2025.

Trang Le et al. Analysis of the Human Protein Atlas Weakly Supervised Single-Cell Classification competition. Nature Methods, 19(10):1221–1229, 2022. doi: 10.1038/s41592-022-01606-z.

Yue Qin et al. A multi-scale map of cell structure fusing protein images and interactions. Nature, 600(7889):536–542, 2021. doi: 10.1038/s41586-021-04115-9.

Alex X. Lu, Oren Z. Kraus, Sam Cooper, and Alan M. Moses. Learning unsupervised feature representations for single cell microscopy images with paired cell inpainting. PLOS Computational Biology, 15(9):e1007348, 2019. doi: 10.1371/journal.pcbi.1007348.

Nicolas Bourriez et al. ChAda-ViT: Channel Adaptive Attention for Joint Representation Learning of Heterogeneous Microscopy Images. In 2024 IEEE/CVF Conference on Computer Vision and Pattern Recognition (CVPR), pages 11556–11565, Seattle, WA, USA, 2024. IEEE. doi: 10.1109/CVPR52733.2024.01098.

Oren Kraus et al. Masked Autoencoders for Microscopy are Scalable Learners of Cellular Biology. In 2024 IEEE/CVF Conference on Computer Vision and Pattern Recognition (CVPR), pages 11757–11768, Seattle, WA, USA, 2024. IEEE. doi: 10.1109/CVPR52733.2024.01117.

Chau Pham, Juan C. Caicedo, and Bryan Plummer. ChA-MAEViT: Unifying Channel-Aware Masked Autoencoders and Multi-Channel Vision Transformers for Improved Cross-Channel Learning. In Advances in Neural Information Processing Systems 38, pages 13342–13372, San Diego, California, USA and Mexico City, Mexico, 2025. Neural Information Processing Systems Foundation, Inc. (NeurIPS). doi: 10.52202/085713-0401.

Peize Sun et al. Autoregressive Model Beats Diffusion: Llama for Scalable Image Generation. arXiv preprint arXiv:2406.06525, 2024.

Salvatore Candido et al. Language Modeling Materializes a World Model of Protein Biology. bioRxiv preprint, 2026. doi: 10.64898/2026.06.03.729735.

Zeming Lin et al. Evolutionary-scale prediction of atomic-level protein structure with a language model. Science, 379(6637):1123–1130, 2023. doi: 10.1126/science.ade2574.

Hugo Touvron et al. LLaMA: Open and Efficient Foundation Language Models. arXiv preprint arXiv:2302.13971, 2023.

Hugo Touvron et al. Llama 2: Open Foundation and Fine-Tuned Chat Models. arXiv preprint arXiv:2307.09288, 2023.

Patrick Esser, Robin Rombach, and Björn Ommer. Taming Transformers for High-Resolution Image Synthesis. In 2021 IEEE/CVF Conference on Computer Vision and Pattern Recognition (CVPR), pages 12868–12878, Nashville, TN, USA, 2021. IEEE. doi: 10.1109/CVPR46437.2021.01268.

Jianlin Su et al. RoFormer: Enhanced transformer with Rotary Position Embedding. Neurocomputing, 568:127063, 2024. doi: 10.1016/j.neucom.2023.127063.

Vineet Thumuluri, José Juan Almagro Armenteros, Alexander Rosenberg Johansen, Henrik Nielsen, and Ole Winther. DeepLoc 2.0: Multi-label subcellular localization prediction using protein language models. Nucleic Acids Research, 50(W1):W228–W234, 2022. doi: 10.1093/nar/gkac278.

Nathan H. Cho et al. OpenCell: Endogenous tagging for the cartography of human cellular organization. Science, 375(6585):eabi6983, 2022. doi: 10.1126/science.abi6983.

Martin Heusel, Hubert Ramsauer, Thomas Unterthiner, Bernhard Nessler, and Sepp Hochreiter. GANs Trained by a Two Time-Scale Update Rule Converge to a Local Nash Equilibrium. In Advances in Neural Information Processing Systems, volume 30. Curran Associates, Inc., 2017.

The UniProt Consortium. UniProt: The Universal Protein Knowledgebase in 2025. Nucleic Acids Research, 53(D1):D609–D617, 2025. doi: 10.1093/nar/gkae1010.

Vidit Agrawal et al. CHAMMI-75: Pre-training multi-channel models with heterogeneous microscopy images. In The Fourteenth International Conference on Learning Representations, 2026.

Ilya Loshchilov and Frank Hutter. Decoupled Weight Decay Regularization. In International Conference on Learning Representations, 2019.

